# Morphoelectric Diversity and Specialization of Neuronal Cell Types in the Primate Striatum

**DOI:** 10.64898/2026.02.26.708019

**Authors:** Xiao-Ping Liu, Rachel Dalley, Nelson Johansen, Agata Budzillo, Julian Thijssen, Jeremy A. Miller, Sarah Walling-Bell, Scott Sawchuk, Lauren Alfiler, Julia Andrade, Angela Ayala, Stuard Barta, Kyla Berry, Darren Bertagnolli, Ashwin Bhandiwad, Madeline Bixby, Krista Blake, Krissy Brouner, Trangthanh Cardenas, Tamara Casper, Anish Bhaswanth Chakka, Thomas Chartrand, Scott Daniel, Nicholas Donadio, Nadezhda I. Dotson, Tom Egdorf, Rachel Enstrom, Yuanyuan Fu, Amanda Gary, Jeff Goldy, Melissa Gorham, Kristen Hadley, Alvin Huang, Avery C Hunker, Atlas Jordan, Zoe C. Juneau, Matthew Jungert, Madhav Kannan, Inkar Kapen, Shannon Khem, Megan Koch, Katalin Kocsis, Rana Kutsal, Gabriela Leon, Matt Mallory, Jocelin Malone, Audrey McCutcheon, Medea McGraw, Kamiliam Nasirova, Dakota Newman, Lindsay Ng, Ximena Opitz-Araya, Alana Oyama, Elliot Phillips, Christina Alice Pom, Lydia Potekhina, Ramkumar Rajanbabu, Shea T. Ransford, Ingrid Redford, Christine Rimorin, Dana Rocha, Augustin Ruiz, Naz Taskin, Michael Tieu, Éva Tóth, Jessica Trinh, Marshall VanNess, Sara Vargas, Maria Camila Vergara, Morgan Wirthlin, Jeanelle Ariza, Trygve Bakken, Nick Dee, Nikolai Dembrow, Song-Lin Ding, Chris English, Samantha D. Hastings, Tim Jarsky, Lauren Kruse, Scott F. Owen, Kimberly A. Smith, Jack Waters, Shenqin Yao, Gábor Tamás, Boudewijn Lelieveldt, Nathan W. Gouwens, Hongkui Zeng, Staci A. Sorensen, Brian R. Lee, Jonathan T. Ting, Ed S. Lein, Brian E. Kalmbach

## Abstract

The basal ganglia are evolutionary ancient subcortical nuclei that form interconnected loops with the neocortex and limbic system to regulate movement, learning, habit formation, emotion, and motivation. Their dysfunction contributes to major neurological and psychiatric disorders, yet most cellular-level insights derive from rodent studies, leaving knowledge gaps in humans and translationally relevant primate species. To address this, we generated multi-modal Patch-seq data linking transcriptomic identity with morphological and electrophysiological properties in macaque striatum, the input nucleus of the basal ganglia. We found underappreciated diversity among medium spiny neurons, including non-canonical types, and variation aligned with functional gradients. Interneurons also exhibited spatial variation and even greater morphoelectric diversity, highlighting their functional modularity. Despite broad evolutionary conservation, we identified primate-specific features and key differences from rodent striatal neurons. By integrating molecular classification with cellular properties that shape network function, our findings provide insights into the functional organization of the primate striatum.

## INTRODUCTION

The basal ganglia are an evolutionarily ancient network of interconnected subcortical nuclei that work in concert with the neocortex and limbic structures through reciprocally connected loops to coordinate volitional movement, motor action selection, learning, habit formation, emotion, cognition, and motivation ^1–4^. Their importance to normal brain function is reflected in their involvement in numerous debilitating neurological and psychiatric disorders including Huntington’s disease, Parkinson’s diseases, depression, and addiction ^5–7,8,9,10^. Remarkably, the core features of basal ganglia circuitry are well conserved across vertebrates, with key aspects of cellular architecture, connectivity and function evident in phylogenetically distant orders like lampreys and primates ^11^. However, with few exceptions ^12^, our knowledge of intrinsic physiology and cellular morphology in primates remains limited, leaving significant gaps in our understanding of how these neuronal features that constrain network function vary across species. Gaining such insights on conservation and divergence is key to uncovering putative primate-specific specializations and more effectively advancing translational research.

The striatum, the primary input nucleus of the basal ganglia, exhibits substantial functional diversity along its dorsal/ventral and anterior/posterior axes, reflecting topographically organized inputs from functionally distinct cortical and thalamic regions ^13,14^. Decades of research — primarily in rodents — have revealed a panoply of principal neurons and interneurons within the striatum ^15–17^. Principal neurons, comprising approximately 95% of striatal neurons, are traditionally classified into two major types of medium spiny neurons (MSNs) based on their projection targets: direct-pathway MSNs projecting to the substantia nigra pars reticulata (SNr) / globus pallidus internus (GPi), and indirect-pathway MSNs projecting via the globus pallidus externus (GPe) and subthalamic nucleus. These pathways are correlated with expression of D1 (direct pathway) or D2 (indirect pathway) dopamine receptors. Direct pathway MSN activation generally promotes movement, whereas indirect pathway MSN activation inhibits it ^18^. However, both populations are recruited during movement initiation, suggesting more nuanced roles than simple opposition^19^. Loss of dopaminergic input to these neurons contributes to the severe motor symptoms of Parkinson’s disease, underscoring their clinical relevance ^20,21^. Interspersed among MSNs are diverse interneurons that modulate the response of MSNs to key afferent pathways, including neocortical, thalamic and dopaminergic inputs from the substantia nigra pars compacta (SNc) and ventral tegmental area (VTA) ^16^. In rodents each of these broad neuronal classes exhibits further variation across anatomical and histochemical subdivisions of the striatum ^5,22,23^. How and to what degree these organizational principles generalize to primates remains unclear.

One fruitful approach to comparing cellular diversity across species has been to leverage single cell transcriptomic data to generate cell type taxonomies in multiple organisms ^24–27^. Aligning these taxonomies across species based on similarity in gene expression reveals cell types in rarely studied orders, like primates, that are homologous to cell types in well studied orders, like rodents. These comparative approaches have uncovered an underappreciated level of cellular diversity within each species, along with notable cross-species differences in the relative abundance of key striatal cell types and the emergence of primate-specific innovations ^23,26,28–32^. However, while transcriptomic data creates a high-resolution map of neuronal types, it alone does not directly reveal cell type-specific or cross-species differences in functional properties, like intrinsic electrophysiology and cellular morphology. Moreover, it lacks validation from non-transcriptomic modalities, which is essential for confirming and refining cell type definitions ^33–35^.

To bridge these gaps in knowledge, we collected multi-modal Patch-seq ^36–38^ data — morphology, electrophysiology, and transcriptomics — from single neurons in the macaque striatum. This approach allowed us to annotate BRAIN Initiative Cell Atlas Network (BICAN) taxonomies with functionally-related properties, enabling comparisons to homologous mouse cell types and existing literature ^30,39^. These data are freely available to the scientific community through open-access data repositories and can be visualized and analyzed through custom tools in the Cytosplore viewer (Figure S1; https://viewer.cytosplore.org/).

## RESULTS

### Macaque striatum Patch-seq pipeline

To study the morphoelectric properties of transcriptomic cell types in the primate striatum, we employed an *ex vivo* brain slice platform that maximizes the use of valuable tissue specimens obtained from animals designated for the Washington National Primate Research Center’s Tissue Distribution program (Figure 1A) ^12,24,40,41^. We conducted Patch-seq experiments in both acute and cultured brain slices and mapped transcriptomic samples to the HMBA Consensus Macaque Basal Ganglia taxonomy ^30^ using computational methods (Figures 1B; STAR Methods). Most analyses presented here were performed at the “Group” level of the taxonomy – just above the terminal cluster level – and the finest level with cross-species generalizability. To facilitate the targeting of rare cell types — such as cholinergic interneurons, which comprise approximately 1% of striatal neurons ^42^ — we used enhancer-driven adeno-associated viruses (AAVs) ^43^ to target specific neuron types based on fluorescent reporter transgene expression in culture. Additionally, we used photo-documentation of specimen dissections and slices to align Patch-seq samples to a three-dimensional macaque reference atlas, enabling anatomical localization and analysis of spatial gradients in physiology and morphology (Figure 1C).

**Figure 1.**
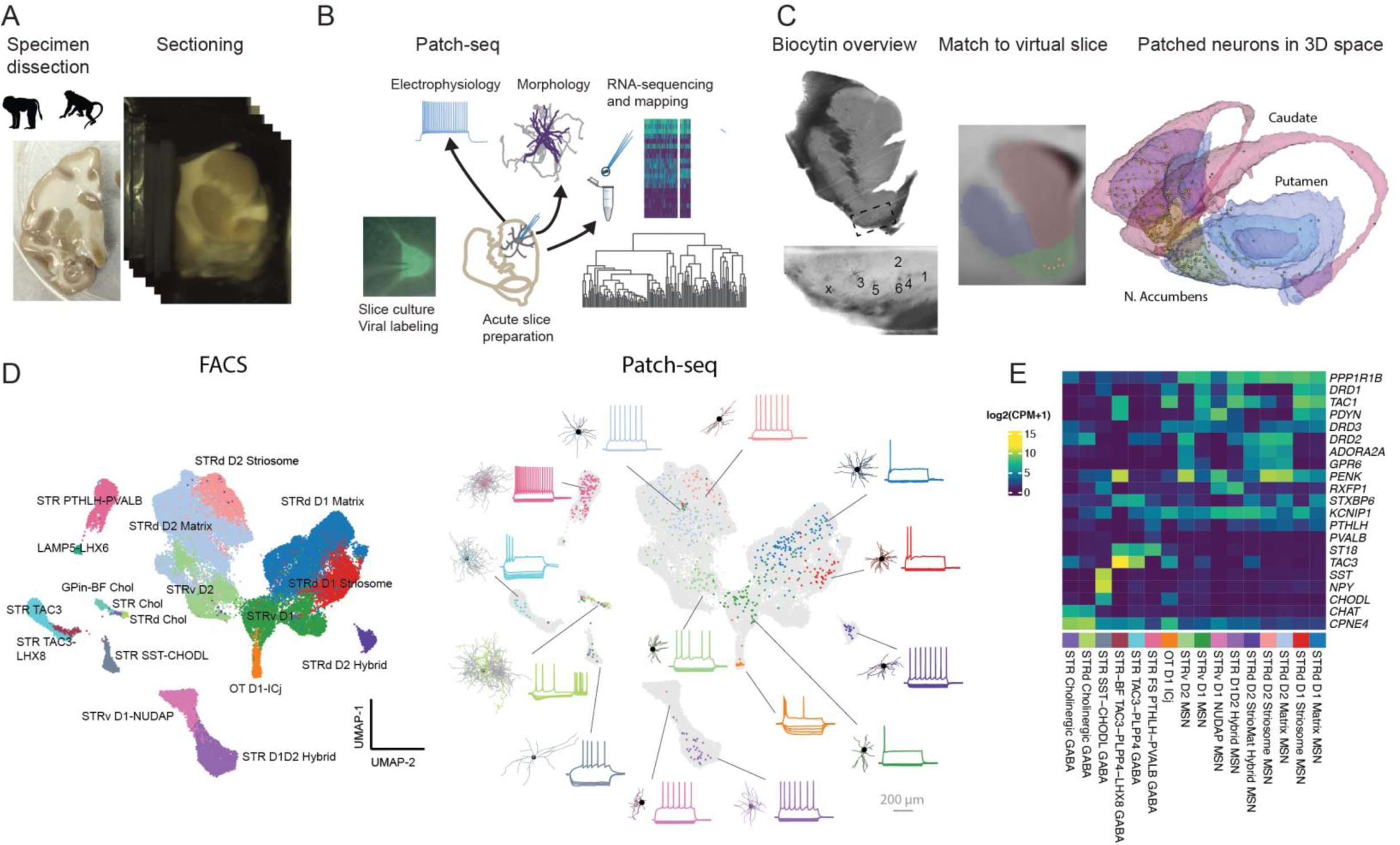
Patch-seq reveals the morphoelectric diversity of transcriptomic cell types in macaque striatum. **A)** Schematic of the experimental workflow. We prepared 300-350 µm thick brain slices from striatal blocks dissected from macaque and squirrel monkey brain tissue obtained from a tissue distribution program. **B)** To maximize tissue yield, Patch-seq experiments were performed in both acute and cultured brain slices. During patch clamp recordings, we assessed intrinsic neuronal properties using a standardized stimulus set and filled neurons with biocytin for subsequent dendritic/axonal reconstructions. The nucleus was then extracted from the recorded neuron, RNA-sequencing was performed, and the resulting transcriptome was mapped to our macaque basal ganglia reference taxonomy. Viral genetic tools were used in slice cultures to aid in targeting relatively rare types, like cholinergic interneurons.**C)** We used photo documentation obtained during slice preparation and biocytin histochemistry to pin samples in a 3D reference atlas. **D)** Integrated UMAP showing reference taxonomy samples (left) and Patch-seq samples (right). Example electrophysiology sweeps and reconstructions from Patch-seq experiments are annotated in the Patch-seq view of the UMAP. **E)** Mean marker gene expression in Patch-seq samples across striatal transcriptomic groups.

Following quality control, our dataset included 716 Patch-seq samples, of which 652 passed electrophysiological benchmarks and 142 exhibited sufficient biocytin fills for dendritic and in some cases axonal reconstructions (Figure S2). These neurons mapped to various transcriptomically-defined MSN and interneuron types (Figures 1D,E). Although sampling was skewed towards MSNs, consistent with their relative abundance ^30^, we sampled from all 17 confirmed striatal neuronal groups ^39^ and achieved a sample size of at least ten in a majority of the major groups (Figure S2). To visualize the transcriptomic identity of Patch-seq samples relative to the reference taxonomy, we integrated gene expression data from Patch-seq samples (SMART-Seq v4) and FACS-sorted samples used to build the taxonomy (10x snRNA-seq) and performed uniform manifold approximation and projection (UMAP; Figure 1D). Patch-seq samples clustered with their corresponding groups in the 10x reference dataset, underscoring the robustness of our mapping approach. Furthermore, key marker gene expression patterns were consistent between Patch-seq and 10x snRNA-seq datasets (Figure 1E), supporting the fidelity of transcriptomic subclass assignments. Importantly, electrophysiological properties in acute and cultured slices were generally similar and differences did not exceed the variation between transcriptomic groups (Figure S2).

### Medium Spiny neurons and related types

Medium spiny neurons (MSNs) and MSN-like neurons showed considerable, largely continuous, transcriptomic heterogeneity (Figure 1D) ^30^. To test whether this molecular diversity was reflected in neuronal properties that may contribute to functional specialization, we characterized the electrophysiological profile and dendritic morphology of individual neurons (Figure 2A). The dendritic morphology of all transcriptomic groups was broadly consistent with the known morphology of MSNs in rodents ^44–46^ and consisted of multiple spiny dendrites radiating from the soma (Figure S3). We measured subthreshold and suprathreshold membrane properties in response to standardized current injection protocols (STAR Methods) and visualized the resulting features in two-dimensional space using UMAP. Neurons with similar transcriptomic identities tended to occupy overlapping regions of electrophysiological UMAP, but no discrete clusters emerged (Figure 2B). This suggests that electrophysiological properties vary systematically across MSN and MSN-like cell types, but in a continuous manner rather than forming sharply delineated groups. Consistent with this interpretation, a classifier trained to predict transcriptomic identity (for groups n ≥ 5) based solely on physiological features achieved performance that was above chance but with limited absolute accuracy (Figure S4). A notable exception was the STR D1D2 Hybrid group, which displayed distinctive physiological characteristics relative to other MSN types.

**Figure 2.**
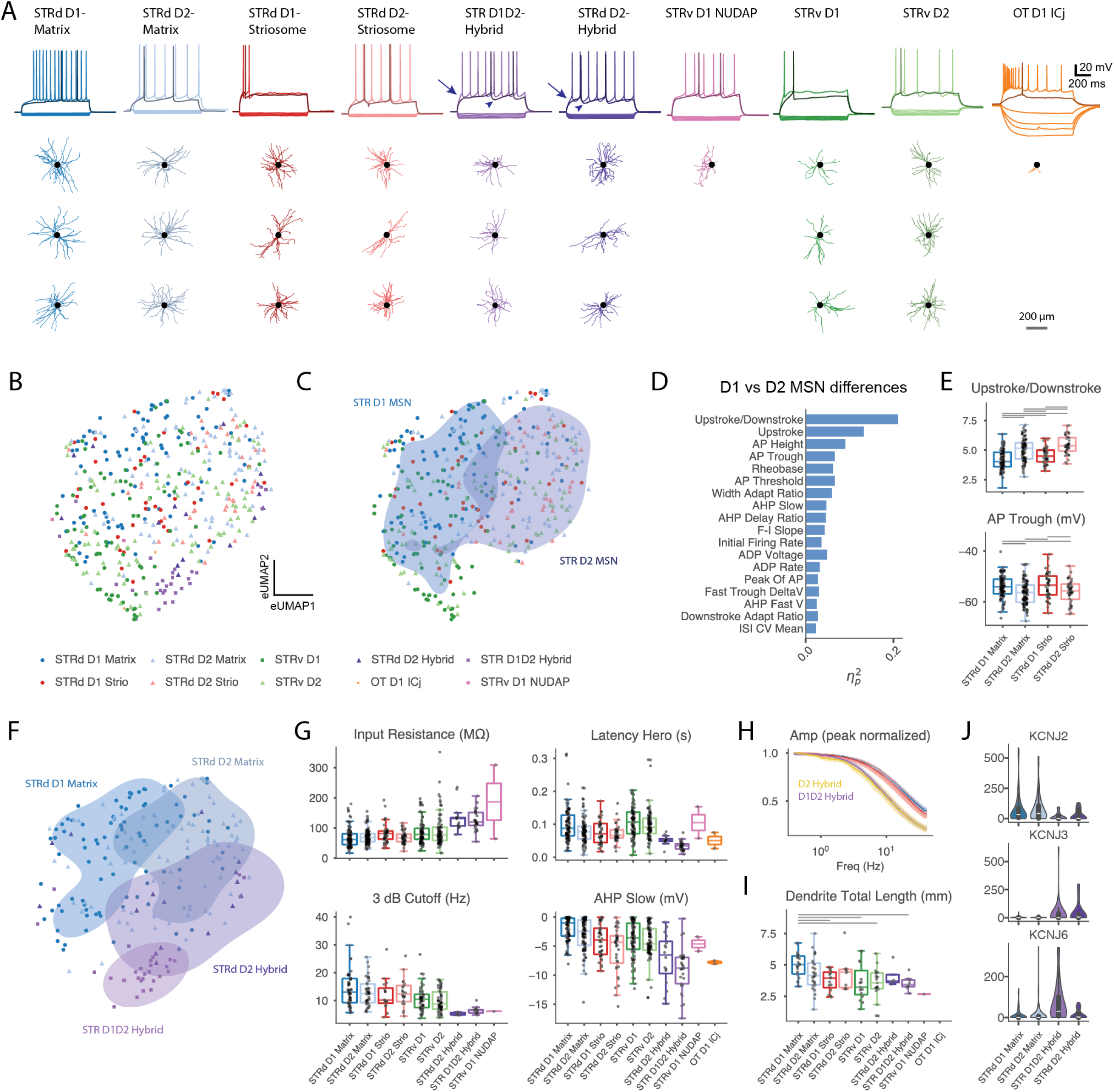
Systematic morphoelectric variation between transcriptomically-defined MSN types in the macaque striatum. **A)** Representative dendritic morphologies and voltage responses to 1s step current injections organized by transcriptomic cell type identity. Rheobase sweep (darker) and +30-40 pA above rheobase (except OT D1 ICj at +15 pA) are shown, as well as standardized hyperpolarizing sweeps (most hyperpolarizing step is –110 pA, except for OT D1 ICj where most hyperpolarizing step is –20 pA due to high input resistance). Arrows and arrowheads illustrate shorter spike latencies at suprathreshold levels and more prominent medium afterhyperpolarizations, respectively, in D1D2-Hybrid and D2-StrioMat Hybrids. **B)** UMAP visualization of electrophysiology features in CN LGE GABA types, color-coded by MSN group; symbols reference subclass (e.g., D1 MSN in circles, D2 MSN in triangles). **C)** UMAP subset showing only D1 and D2 MSNs with kernel density estimate contours at 50% of peak density overlayed for the STR D1 MSN and STR D2 MSN subclasses. **D)** Partial effect sizes for electrophysiological features distinguishing D1 from D2 MSNs for both matrix and striosome compartments (D1/D2 effect – two-way ANOVA with D1/D2 and matrix/striosome as factors). Sorted by descending FDR corrected *p*-values, which can be found in Figure S4. **E)** Example features that distinguish D1 and D2 MSNs: action potential upstroke/downstroke ratio (top) and post-spike voltage trough (bottom) for main dorsal striatum MSN groups. **F)** UMAP from panel B, subset to matrix and non-canonical MSN types. **G)** Electrophysiology features that differentiate non-canonical MSN types from other MSN types. OT D1-ICj excluded where unavailable or far outside y-range. **H)** Mean normalized impedance amplitude profiles derived from chirp stimulus responses. D1 Matrix (n= 43), D2 Matrix (n = 32), D1 Striosome (n = 14), D2 Striosome (n = 15), D1D2 Hybrid (n = 10), D2 StrioMat Hybrid (n =6, color adjusted for visibility). Error bands indicate the standard error of the mean (SEM). **I)** Total dendritic length across MSN groups. **J)** Ion channel gene expression from Patch-seq experiments from main matrix MSN types and non-canonical types; y axis is in counts per million mapped reads. In panels **E)** and **I)**, horizontal bars mark comparisons significant at α = 0.05 after FDR correction. Only groups with ≥3 samples were tested.

The MSN transcriptomic groups could be considered either by their predominant localization to particular histochemical or anatomical divisions within the striatum, or in terms of their identity as unique transcriptomic types that may be broadly intermixed ^29,30,39^. We asked whether the physiological variation we observed relates to these divisions by grouping transcriptomic cell types based on their anatomical localization and/or their membership in distinctive groups and testing for morphological and physiological differences. These groups, which we expound upon below, are: 1) Dorsal striatum matrix/striosome cell types (STRd D1 Matrix, STRd D2 Matrix, STRd D1 Striosome, STRd D2 Striosome), 2) non-canonical cell types (STRd D2 Hybrid, STR D1D2 Hybrid, STRv D1 NUDAP) and 3) ventral cell types (STRv D1, STRv D2, OT D1 ICj). STRv D1 NUDAP is both a ventral MSN-like type spatially and a non-canonical type transcriptomically; we discuss it with the non-canonical types.

#### D1 versus D2 and matrix versus striosome MSNs

In addition to the classic organization comprised of the D1-direct and D2-indirect pathways, there are histochemically distinct compartments called striosomes (or patches), which form a three-dimensional labyrinth within the surrounding matrix ^22,47^. Although both compartments contain D1 and D2 MSNs that exert opposing effects on behavior, their roles differ. Matrix D1 MSN activation promotes movement and matrix D2 MSN activation suppresses it. Conversely, and perhaps counterintuitively, striosomal D1 MSN activation inhibits movement and striosomal D2 MSN activation facilitates it^48^. These functional differences are thought to reflect distinct connectivity. Striosomes preferentially receive limbic input and project to dopaminergic neurons in the substantia nigra pars compacta (SNc) and distinct regions of the GPe, while the matrix is primarily innervated by cortical afferents and projects to the SNr/GPi and GPe ^5,49^. Beyond connectivity, rodent studies demonstrate that D2 MSNs are more excitable than D1 MSNs, possibly reflecting their lower dendritic surface area ^46,50,51^. Striosomal D1 MSNs in rodents are also more excitable than their matrix counterparts, but the electrophysiological properties of striosomal D2 MSNs are largely unexplored ^52^. A recent study in macaque putamen found that the D1 MSN vs. D2 MSN excitability difference is more subtle as compared to rodents ^12^, and matrix vs striosome distinctions in morphology and physiology are not well characterized.

To address this knowledge gap, we performed a two-way ANOVA on electrophysiological features with main MSN type (D1 vs D2) and compartment (matrix vs striosome) as factors. This revealed several electrophysiological features that differentiated D1 from D2 MSNs across both domains (Figures 2D and S4). For example, as reported previously ^12^, D2 MSNs displayed less symmetrical spikes (higher upstroke/downstroke ratio), a lower action potential threshold and more negative after spike voltage troughs (Figures 2E and S4). Notably, many D1 vs. D2 MSN differences previously reported in rodents were more subtle, often not reaching statistical significance, but directionally consistent in primate MSNs (Figures 2G and S4). These include lower rheobase, higher input resistance (R_in_, p = 0.91), and lower membrane time constant (tau, *p* = 0.90) in D2 MSNs. Matrix vs. striosome differences generally had smaller effect sizes, although some clear effects emerged such as striosomal MSNs having larger slow component of afterhyperpolarizations following spikes (“AHP Slow”) on top of the D1 vs. D2 MSN effect for that feature (Figure S4). Consistent with these observations, D1 and D2 MSNs occupied different parts of electrophysiological UMAP space (Figure 2B-C).

The dendritic morphology was also very similar between matrix and striosome MSNs, but D1 matrix MSNs exhibited the greatest total dendritic length (Figures 2A, I and S4). Some striosomal MSNs had dendrites oriented along a single axis rather than radially, (Figures 2A and S3) consistent with observations in rodents that striosomal dendrites are confined within their compartment boundaries ^53^.

#### Non-canonical medium spiny neuron types

In addition to the canonical dorsal striatum MSN types, we studied three atypical populations that together comprise a sizeable minority of the overall MSN and MSN-like pool: STR D1/D2 hybrid, STRv D1 NUDAP, and STRd D2 Hybrid neurons ^30,39^. The STR D1/D2 Hybrid neurons occupy a distinct “third island” region in transcriptomic UMAP space, clearly separated from canonical MSN clusters (Figure 1D). These neurons have been previously described under various names — including D1-Pcdh8 ^28^, eccentric SPNs ^54^, and D1H ^31^. D1 NUDAP neurons also occupy the “third island” and correspond to a previously molecularly characterized MSN-like population that forms spatial clusters in <u>n</u>eurochemically <u>u</u>nique <u>d</u>omains in the nucleus <u>a</u>ccumbens and ventral <u>p</u>utamen (NUDAP)^29^. The corresponding mouse type, STR-PAL Chst9 Gaba, is assigned to the intercalated amygdalar nucleus^55^ and intercalated cells have been inferred to share a developmental origin with eccentric MSNs^56^, suggesting that MSN-like terminal groups extend beyond the striatum. Our sample of STRv D1 NUDAP MSNs was very small, so we report those data with minimal interpretation. STRd D2 Hybrid neurons represent a novel population of MSNs that co-expresses marker genes of both striosome and matrix compartments ^30^. Collectively, we refer to these three groups as non-canonical MSNs, whose presence suggests a more nuanced cellular architecture than the classic binary framework. Notably, these types may constitute a larger proportion of the total MSN population in primates compared to bats and mice ^30,57^, highlighting their potential importance in the primate striatal circuit.

We asked whether these populations exhibit distinctive electrophysiological and morphological properties, consistent with their unique transcriptomic profiles. Indeed, on a UMAP visualization of electrophysiological properties, STR D1D2 Hybrid MSNs were clustered in a clearly delineated region, consistent with their relative ease of classification, while STRd D2 Hybrid MSNs were intermediate between STR D1D2 Hybrid and STRd D2 MSN regions (Figures 2B,F and S4). Although many electrophysiological features overlapped with canonical MSN types, key differences emerged (Figure 2G). Non-canonical MSNs had higher input resistance and lower rheobase, indicating that less input is required to elicit spiking (Figures 2G and S4). Their subthreshold voltage responses to time-varying inputs also differed, showing stronger low-pass filtering as quantified by a lower 3dB cutoff frequency (Figure 2G, H). These distinctions were associated with differences in the expression of genes encoding several types of inward rectifier K+ channels (Kir), which contribute to the hyperpolarized resting potential and low input resistance of canonical MSNs ^58,59^ (Figure 2J). Non-canonical types expressed lower levels of *KCNJ2* (Kir2.1), which may contribute to their relatively higher input resistance. In contrast, non-canonical MSNs expressed higher levels of G-protein coupled Kirs (*KCNJ3 and KCNJ6; GIRK1/*Kir3.1 *and GIRK2/*Kir3.2), suggesting enhanced neuromodulatory control through receptors that act on GIRK channels like opioid, GABA_B_ and dopamine D2-like receptors ^60–62^.

Among non-canonical types, STR D1D2 Hybrids were particularly distinctive (Figures 2F, S4). These neurons showed a prominent slower component of the afterhyperpolarization potential (Figure 2A, G), along with adaptive changes in spike threshold (Fig S4). These differences may reflect elevated T-type Ca²⁺ channel expression (CACNA1G/CACNA1H) in non-canonical MSNs, which can couple to Ca²⁺-dependent K⁺ channels to produce more pronounced AHPs, and/or greater *KCNT2* (Na⁺-activated K⁺) expression ^63,64^ (Figure S4). Such adaptive properties may account for the generally lower maximum firing rates we observed in STR D1D2 Hybrids, despite their higher input resistance relative to canonical MSNs. We confirmed that prominent contrasts such as in the slow component of the afterhyperpolarization or the latency to first spike were not confounded by culture condition or the anatomical location of the sample (Figure S4).

#### Ventral MSNs

The dorsal/ventral axis represents another important anatomical and functional division of the striatum. The dorsal striatum (caudate/putamen) is primarily involved in movement planning and sensory/motor integration, whereas the ventral striatum —including the nucleus accumbens (NAc) and ventral portions of the caudate and putamen—processes limbic input and contributes to emotional learning and addiction ^65,66,2^. Transcriptomically, the dorsal-ventral axis appears to form a continuum within both D1 and D2 subclasses, with the ventral populations annotated as STRv D1 and STRv D2 MSNs (Figure 1D; companion papers ^30,39^). Notably, STRd Matrix MSNs predominantly localized to the dorsal striatum, whereas STRv MSNs occupy the NAc and ventral regions of the caudate and putamen (Figure S7). In our reference taxonomy and companion spatial transcriptomics work^30,39^, the NAc core and shell are not reliably demarcated by markers across primate species, and cell types do not segregate clearly along this boundary, limiting our resolution at this level.

Comparing morphoelectric properties between dorsal and ventral MSN types, we observed systematic variation consistent with these anatomical and functional delineations. Differences were most pronounced in the D1 MSN subclass — for example, between STRd D1 Matrix and STRv D1 — and are evident in the electrophysiological UMAP (Figure 3A), the MSN classifier (Figure S4), and dedicated spatial analysis section (Figure 6). STRv D1 MSNs, particularly those located in the NAc, could exhibit broad action potentials with shoulder-like inflections during repolarization (Figure 3B,C). Notably, action potential width was inversely correlated with *KCND2* expression, implicating Kv4.2 channel expression in shaping kinetics, consistent with findings in rodent neocortical neurons^67^ (Figure 3E, Spearman’s correlation coefficient = −0.25, p-val = 1.3e-8). Additionally, the action potential upstroke was slower in ventral MSNs and the slow component of the afterhyperpolarization potential was larger, with the latter more tightly linked to transcriptomic identity than anatomical location (Figure 3C). Ventral MSNs had lower total dendritic length than STRd D1 MSNs, and STRv D2 MSNs possessed fewer dendritic branches (Figures 3D; S3).

**Figure 3.**
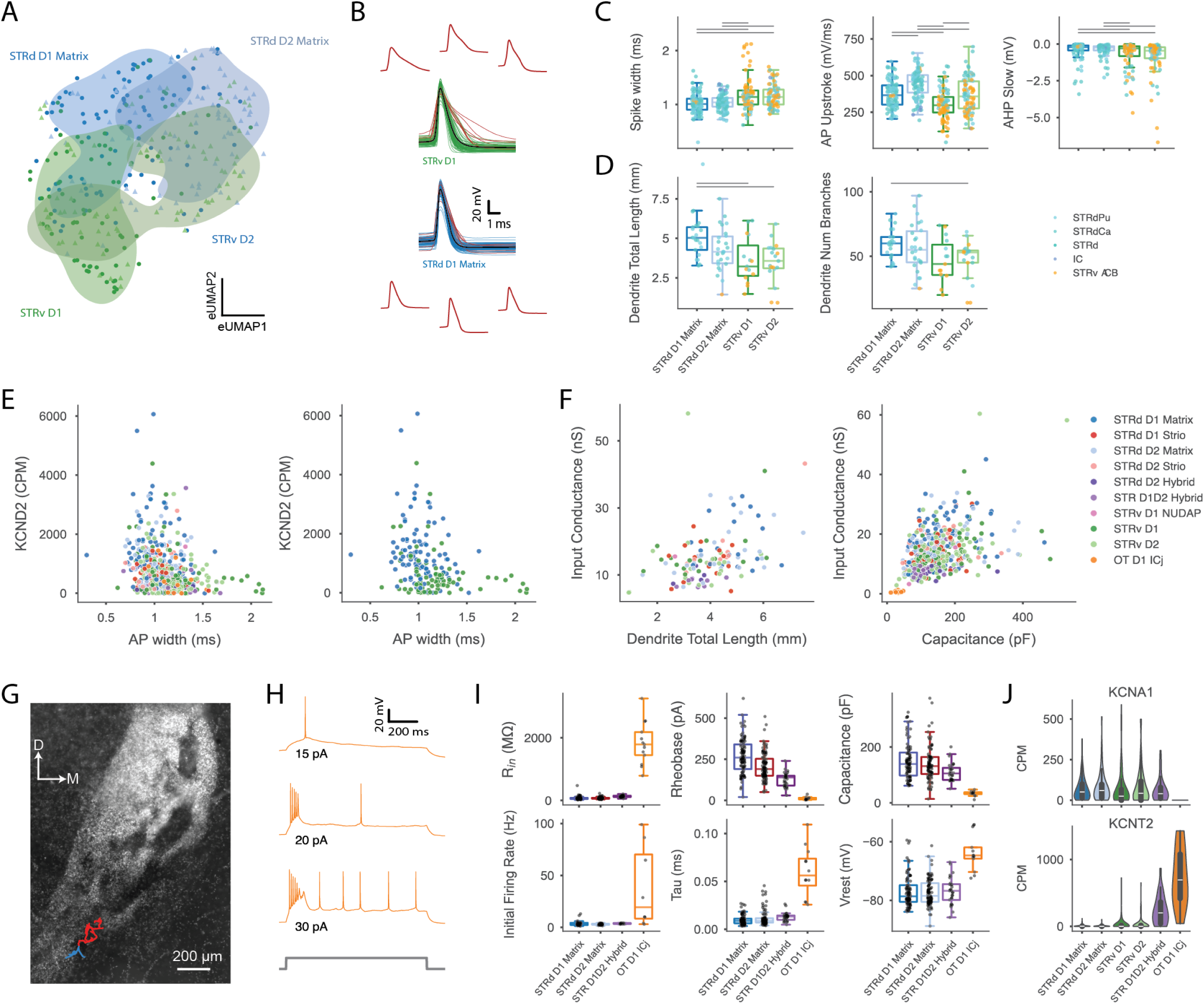
Unique properties of ventral LGE-derived neuron types. **A)** Same electrophysiology UMAP as in Fig. 2B, subsetted to dorsal matrix and ventral MSN groups. **B)** Overlays of peak-aligned action potential waveforms for STRv D1 vs. STRd D1 Matrix MSNs. The seven widest spikes in each group are highlighted in red, and three of those are shown separately. **C)** Action potential features that differ between ventral and dorsal MSNs, with the anatomical location indicated by color (bars denote FDR corrected p-value < 0.05). **D)** Total dendritic length and number of branches for dorsal matrix and ventral MSN groups. **E)** *KCND2*, which encodes Kv4.2, plotted as a function of AP width across all LGE groups (left) and just the STRd D1 Matrix and STRv D1 types (right). **F)** Input conductance (1 / input resistance) plotted against features relating to total dendritic surface area: total dendritic length (left) and capacitance (right). **G)** Morphological reconstruction of an OT D1 ICj neuron superimposed upon DAPI stain to reveal the high-density island of granule cells. Axon in red, dendrites in blue. **H)** Example of bursting behavior in an OT D1 ICj cell in response to step current injection. **I)** Electrophysiological features that distinguished OT D1 ICj from other LGE-derived neurons, including input resistance (R_in_), threshold current (rheobase), capacitance, 1 / First ISI (initial firing rate), membrane time constance (tau), and resting membrane potential (Vrest). **J)** Expression of *KCNA1* (Kv1.1) and *KCNT2* (SLICK) in OT D1 ICj and MSN types.

#### Islands of Calleja Granule Cells

In addition to ventral striatal MSN types, we sampled from neurons within the Islands of Calleja (ICj), dopamine D3 receptor-rich, cell body-dense regions embedded within the olfactory tubercle (OT). Optogenetic activation of these neurons triggers grooming behavior in mice, while inhibition halts ongoing grooming^68^. Additionally, activation of these neurons is rewarding while ablation or inhibition produces depressive symptoms ^69^. Accordingly, they are thought to be the site of action of anti-psychotic and anti-depressant drugs acting on the dopamine D3 receptor ^70^, and have been hypothesized to play a role in dopamine dysfunction in schizophrenia ^71^. Despite their clinical relevance, the cellular properties of ICj granule neurons remain poorly characterized, especially in primates.

These OT D1 ICj neurons were considered an MSN-like cell type in He et al. (2021), share the CN LGE GABA class and markers of the lateral ganglionic eminence (LGE) with MSN types in our taxonomy, and can appear contiguous with ventral D1 MSN’s on a transcriptomic UMAP (Figure 1D). However, in contrast to MSNs, they have a granule cell morphology and likely do not project beyond the olfactory tubercle ^68,70^. Although several electrophysiological properties of our Patch-seq OT D1 ICj neurons fell within the MSN range (Figures 2G, S4), other features were strikingly different between these cell types. While MSNs are known for their low input resistances, our primate OT D1 ICj neurons had an input resistance more than an order of magnitude higher, along with low membrane capacitance, long time constant and a low rheobase (Figure 3I), similar to reports in rodent ICj neurons ^68–70,72^. The higher input resistance can largely be explained by reduced dendritic membrane area (Figure 3F), for which capacitance serves as a proxy, since conductance scales with membrane area. These interconnected features are consistent with a cell type with small and simple morphologies. Indeed, our sole reconstructed macaque OT D1 ICj neuron had only a few short dendrites and an axonal process emanating in the opposite direction (Figure 3G), similar to mouse ICj granule cells ^70^.

Surprisingly, unlike previous reports describing rodent ICj granule cells as regular-spiking neurons ^68–70,72^, we observed a spectrum of firing patterns in macaque OT D1 ICj neurons ranging from regular firing to onset bursting (Figure 3H). In fact, a majority (13/15) of macaque OT D1 ICj neurons exhibited some indication of bursting behavior, most typically high frequency bursts riding on a depolarized envelope followed by low-rate regular firing (e.g., Figure 3H). Although bursting behavior in some neurons developed at current injections higher than rheobase, this bursting propensity could nonetheless be captured by the inverse of the first interspike interval at rheobase (Figure 3I, initial firing rate). Notable ion channel expression differences include absence of *KCNA1* (K_V_1.1) and elevated *KCNT2* (K_Na_/Slick) expression in OT D1 ICj neurons (Figure 3J), encoding for a low-threshold potassium channel that dampens near threshold excitability and a sodium-activated potassium channel that could reduce excitability following burst firing, respectively.

### Interneurons

In contrast to MSN types, the morphoelectric properties of interneurons were distinctive (Figures 4A, S5). Morphologically, total dendritic length and number of branches varied greatly between interneuron groups (Figures 4A, E, and S5). In UMAP views of electrophysiological properties interneurons formed clearly separated clusters (Figure 4B) and transcriptomic identity could be predicted based solely on electrophysiology with high accuracy (Figure 4C). Similarly, ANOVA revealed that both subthreshold and suprathreshold membrane properties varied substantially across interneuron groups, with differences generally exceeding those seen in MSNs (Figure 4D,E, and S5). Interneurons were distinguished by subthreshold membrane properties, including membrane time constant, resting membrane potential, input resistance, and relative size of hyperpolarization-induced sag. Likewise, suprathreshold features like action potential width and after spike potentials, also distinguished interneuron groups. These observations highlight the pronounced morphoelectric diversity of striatal interneurons.

**Figure 4.**
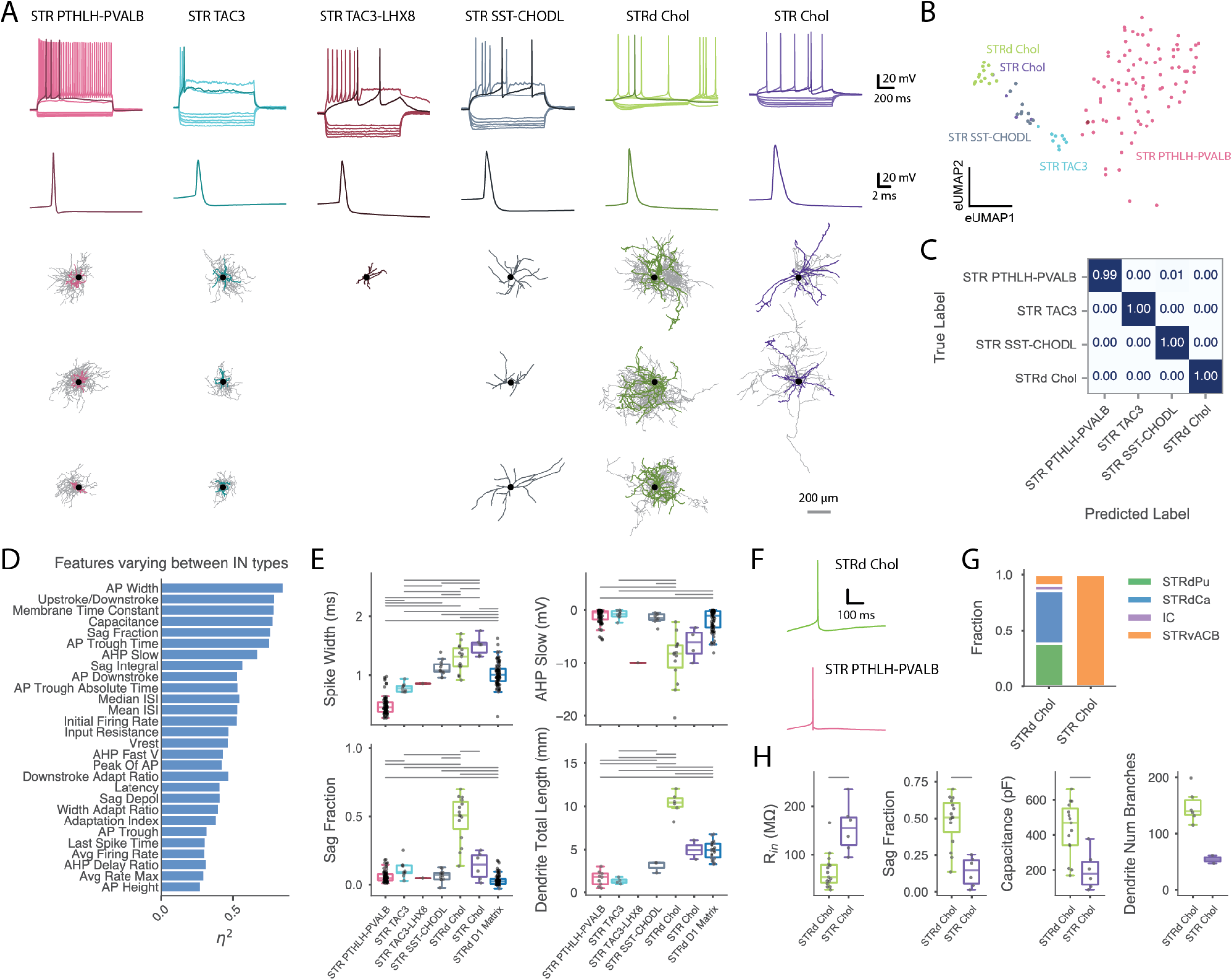
Signature morphoelectric properties of striatal interneuron types. **A)** Representative dendritic (color) and axonal (gray, where available) morphologies and voltage responses to 1s step current injections. Rheobase sweep (darker) and +30-40 pA above rheobase are shown, as well as standardized hyperpolarizing sweeps (most hyperpolarizing step is –110 pA). **B)** UMAP visualization of electrophysiology features in CN MGE GABA types, color-coded by transcriptomic Group. **C)** Normalized confusion matrix for logistic regression classifier trained to categorize interneurons; groups with less than 5 samples with complete feature sets were excluded. **D)** Top features from a one-way ANOVA across interneuron groups (features ranked by p-value; see Figure S5; bars denote FDR corrected p-value < 0.05). **E)** Example electrophysiological and morphological features that distinguish striatal interneurons: action potential half-width (top left), slow component of the afterhyperpolarization (top right), hyperpolarization sag fraction (bottom left) and dendritic total length for interneuron types and D1 Matrix MSNs for comparison. **F)** Representative action potential traces contrasting STRd Chol neurons with STR PTHLH-PVALB neurons, highlighting the pronounced slower component of the AHP in the cholinergic cells. **G)** Stacked bar plot showing the regional distribution of mapped cells, with STRd Chol neurons primarily from dorsal striatum and STR Chol neurons from ventral striatum. **H)** Physiological and morphological features that differ between STRd Chol and STR Chol neurons. Abbreviations: STRdPu, putamen; STRdCa, caudate; IC, internal capsule; STRvACB, nucleus accumbens.

STR PTHLH-PVALB neurons corresponded to the classical fast-spiking interneuron population, characterized by dense radially oriented dendritic and axonal arbors that resembled those of basket cells^16,23,73^. These neurons had the narrowest action potentials, minimal sag, and the fastest time constants (Figures 4E, S5). STRd Chol and STR Chol cholinergic interneurons correspond to tonically active neurons (TANs) or CHAT+ neurons described in rodents, with large somata and extensive axonal branching ^16,73,74^ (Figure 4A). Their electrophysiology was characterized by broad spikes, a prominent slow component of the afterhyperpolarization, and a depolarized membrane potential (Figures 4E, F and S5), which are reminiscent of their rodent counterparts ^75^. STR TAC3 and STR TAC3-LHX8 appear to be dorsal and ventral subtypes of a TAC3+ interneuron population recently described in primates ^26,27,76^. Their dendritic and axonal morphology closely resembled that of STR PTHLH-PVALB neurons (Figure 4A), and their electrophysiological properties were intermediate in spike width and time constant (Figures 4E, S5). STR SST-CHODL neurons resembled SST+/NOS+/NPY+, low-threshold spiking interneurons in rodents, with long, sparsely branched dendrites, high input resistance, minimal sag, and in some neurons spontaneous tonic firing or slow plateau potentials ^16,73^ (Figures 4A,E, S5, S6). Interestingly, mouse striatal Sst-Chodl cells appear related to long-range projecting cortical Sst-Chodl neurons ^23,77–80^. Lastly, we sampled one LAMP5-LHX8 neuron, which expressed marker genes associated with neurogliaform cells, including *NPY* but not *SST*, *DRD1*, *DRD2*, or *CHAT* (Muñoz-Manchado et al., 2018). In rodents, neurogliaform cells, although exceedingly rare, exert powerful inhibition on the striatal circuit ^81,82^. The electrophysiological properties of this cell were consistent with rodent neurogliaform cells and consisted of a regular firing phenotype, moderate spike width and an intermediate input resistance (Figure S6).

#### Distinctive properties of cholinergic interneurons

Cholinergic neurons exhibited a longer AHP than other interneuron types (Figure 4E,F), which peaked at ∼50 ms, within the range of a medium AHP. Note that our analysis feature “AHP Slow” refers to the slow component of the afterhyperpolarization does not specifically imply very slow time courses often referred to as slow AHP. This component is typically caused by small conductance calcium-dependent potassium channels (SK, *KCNN1-3*) or Kv7 (*KCNQ*) channels ^83^ and is significantly reduced in rat striatal cholinergic interneurons by the SK channel blocker apamin (Bennett et al., 2000). In agreement, we detected the expression of KCNQ and *KCNN2/3* genes in the cholinergic groups.

A hallmark of cholinergic striatal interneurons, observed in both rodents and primates, is their tonic firing pattern, upon which transient pauses then rebounds in firing are superimposed in response to motivationally salient stimuli ^84,85^. This tonic activity is generated by intrinsic ionic mechanisms and persists even in the absence of fast synaptic transmission ^74^. Consistent with this, we observed tonic firing in the presence of synaptic blockers in macaque cholinergic interneurons, with 20 out of 26 recordings showing either highly regular or repetitively bursting spontaneous firing in cell-attached and whole-cell configurations (Figure S5). The presence of bursting and regular firing modes was also observed in rodent ^74^ and may relate to the balance between calcium currents and calcium-dependent K^+^ currents ^86^.

There are two groups of striatal cholinergic interneurons in the transcriptomic taxonomy – STRd Chol is mostly localized dorsally, while STR Chol is found ventrally (Figure 4G). Because the ventral localization of STR Chol does not generalize from macaque to all other primate species in the taxonomy, the regional identifier remains “STR” rather than “STRv” ^39^. In rodents, cholinergic neurons are characterized by a prominent hyperpolarization-activated sag mediated by HCN channels ^87^. We found this feature to be pronounced in STRd Chol neurons, but less so in STR Chol neurons (Figures 4E,H and S5), suggesting anatomical or cell type variation in HCN channel expression or function. Indeed, *HCN4* expression – which is associated with slow sag and pacemaking activity ^88^ – was higher in STRd cholinergic neurons (Figure S5). Additionally, input resistance was higher and membrane capacitance lower in STR Chol neurons (Figure 4D,G), suggesting that they may possess less membrane surface area than STRd Chol neurons. Consistent with this, STRd Chol neurons had more dendritic and axonal branches and greater total dendritic length than STRv neurons (Figures 4A, E and S5). Together, these findings reveal morphoelectric distinctions between dorsal and ventral cholinergic interneuron subtypes in the primate striatum, underscoring potential functional specialization aligned with anatomical domains.

#### Distinctions between striatal and neocortical fast spiking interneurons

Striatal PTHLH neurons resemble fast-spiking, basket-cell–like interneurons observed in multiple brain regions, including neocortex ^89^. These neurons are characterized by narrow action potentials, the ability to sustain high firing rates, and their role in entraining local circuits to gamma-band oscillatory activity ^16,73,90,91^. They typically express the calcium-binding protein, parvalbumin, albeit at variable levels ^23,92^, and we collectively refer to them here as fast-spiking (FS) interneurons.

In rodents, neocortical and striatal FS interneurons display distinct electrophysiological properties, demonstrating that interneuron properties can differ markedly across brain areas even among homologous, lineage-related cell types ^23^. To determine whether similar area-specific differences exist in macaques, we collected Patch-seq samples from neocortical PVALB interneurons and compared their properties to striatal PTHLH-PVALB interneurons. In macaques, neocortical and striatal FS interneurons were clearly separable, occupying distinct regions in electrophysiology UMAP space (Figure 5A). As observed in rodents, several intrinsic membrane properties differed between FS interneurons in striatum and cortex, although the specific patterns in macaque diverged from those in mice. Cortical FS interneurons in macaque exhibited narrower action potentials with faster repolarization, higher spike thresholds and deeper fast afterhyperpolarizations compared to striatal FS neurons (Figures 5B and S6). Consistent with these physiological differences, the expression of *KCNC2,* which encodes a Kv3 channel supporting fast action potential kinetics ^93,94^, was higher in cortical FS interneurons than striatal FS interneurons (Figure 5C). HCN channel–dependent properties also varied between areas. Sag was markedly greater in cortical FS interneurons (Figure 5B), whereas in mice sag is largely absent in both regions ^23,95^. Moreover, in macaque cortical FS interneurons, we observed band-pass filtering with a resonance peak around ∼8 Hz in response to time varying inputs (Figure 5D, E). In contrast, striatal FS interneurons exhibited low-pass filtering (Figure 5D, E). Restorative currents, like those mediated by HCN channels, can contribute to subthreshold membrane resonance in a similar frequency band ^96^. The expression of *HCN1,* which encodes a major pore forming subunit of HCN channels ^88^, was higher in cortical FS interneurons (Figure 5C, *HCN2* not significantly different), consistent with their more pronounced HCN-dependent properties. Finally, rheobase was higher and input resistance lower in macaque striatal FS interneurons (Figures 5B and S6), the inverse of the pattern observed in mice, where cortical FS interneurons have lower rheobase and higher input resistance ^23^. Notably, rheobase is substantially higher in mouse striatal FS interneurons than their macaque counterparts (Figure 7D), suggesting these incongruities may reflect genuine cross-species shifts between regions. Together, these findings indicate that macaque striatal FS neurons differ from their cortical counterparts in ways that are distinct from those reported in mice.

**Figure 5.**
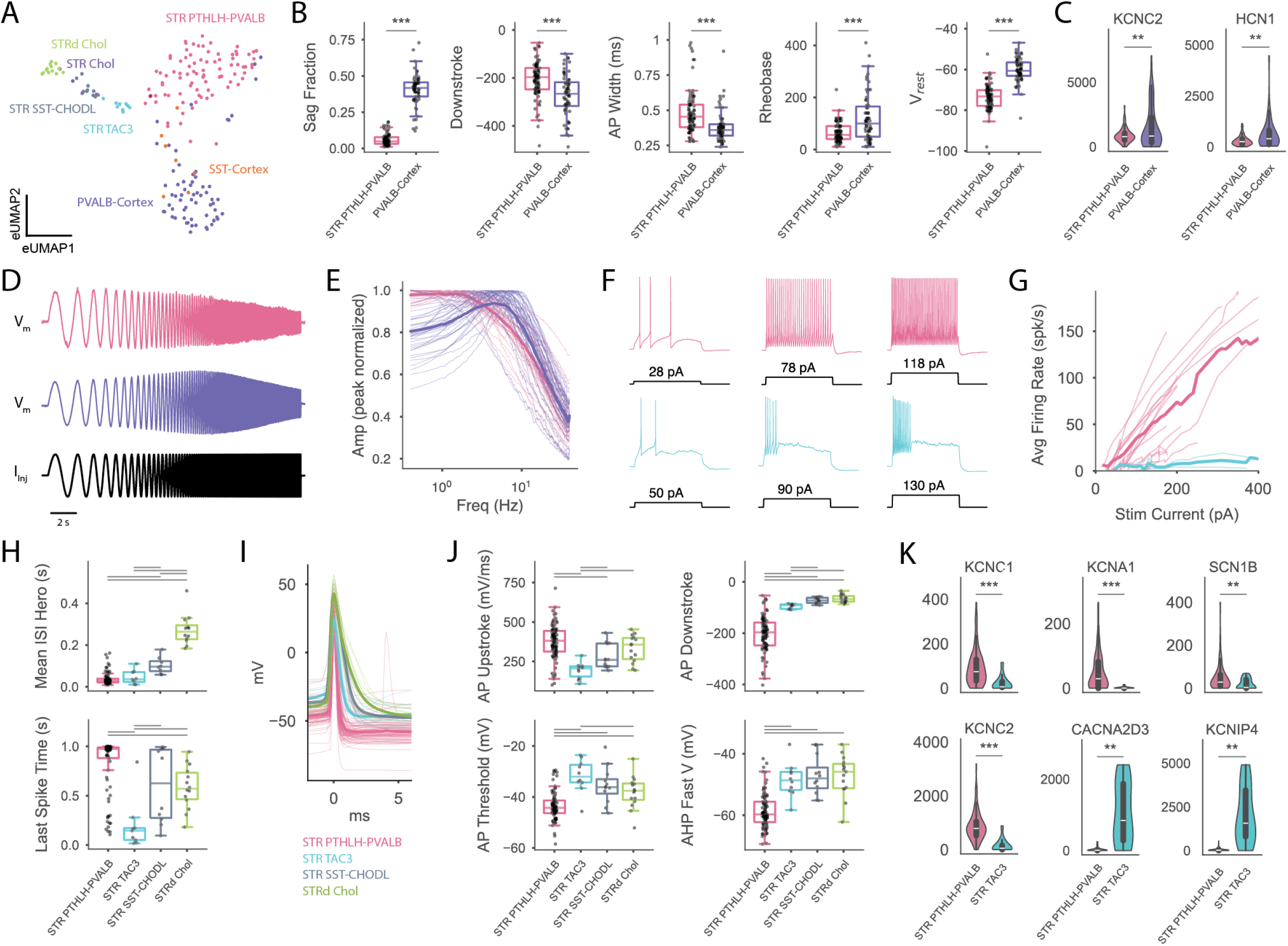
PTHLH interneurons are distinct from cortical fast-spiking interneuron types and TAC3 interneurons. **A)** Integrated UMAP as in Fig. 4B also showing macaque cortical samples labeled with the subclass PVALB or SST. **B)** Compared with cortical FS interneurons, striatal FS interneurons (STR PTHLH-PVALB) had wider APs and slower downstrokes, and very little detectable hyperpolarization sag, among other differences **C)** These electrophysiological differences were consistent with lower expression of KCNC2 and HCN1 in the striatal FS population (units in counts per million). **D)** Example responses of a striatal (top) and cortical (bottom) FS interneuron to a 50 pA sinusoidal frequency-sweep (‘chirp’) current stimulus; voltage axis ranges differ for the two neurons. **E)** Normalized impedance amplitude profiles highlighting a prominent resonant peak in the response of cortical, but not striatal, FS interneurons. Thick lines represent group average. **F)** Representative suprathreshold sweeps show abrupt termination of firing early in STR TAC3 neurons versus sustained firing in STR PTHLH-PVALB neurons. **G)** Average firing rate versus current amplitude for STR PTHLH-PVALB and STR TAC3 neurons. Thick lines indicate group averages and thin lines represent randomly sampled individual cells. **H)** Mean interspike interval and time of last spike during the step computed at “hero” level (30-40 pA above threshold). TAC3 neurons can reach similarly short ISIs at this level but are unable to sustain firing. **I)** Average (thick) and individual (thin) peak-aligned AP waveforms for each major interneuron group. **J)** Action potential properties that differentiate TAC3 and PTLH interneurons. **K)** Differences in ion channel–related gene expression between PTHLH and TAC3 interneurons that may contribute to their distinct intrinsic electrophysiological properties (units in counts per million).

**Figure 6.**
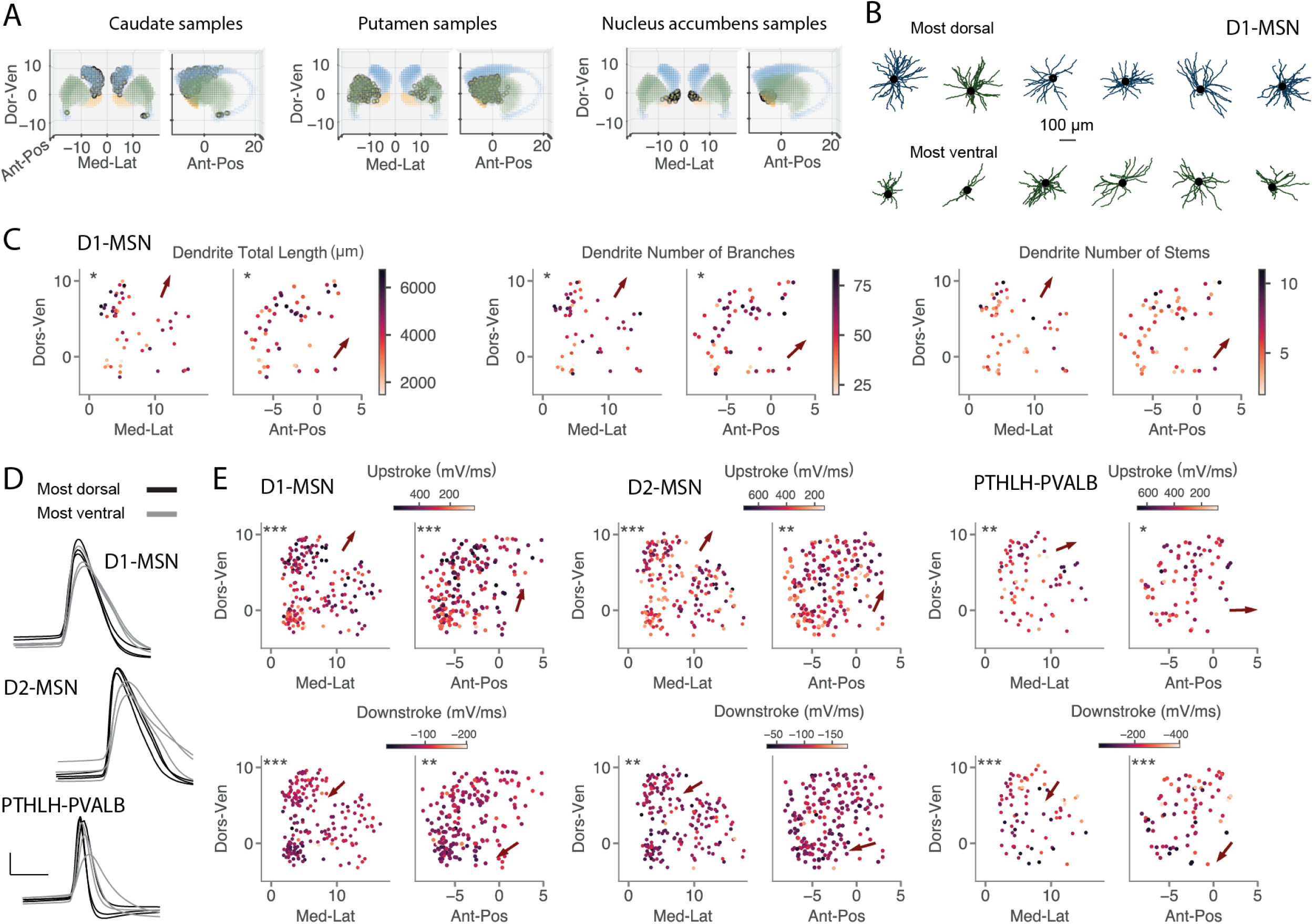
Spatial variation in neuronal properties. **A)** Locations of somas of Patch-seq samples were registered to a macaque anatomical reference (STAR Methods). Panels show the locations of samples assigned to each striatal structure (caudate in *blue,* putamen in *green*, n. accumbens in *orange*). **B)** Example reconstructions for the extremes of the spatial gradient, roughly corresponding to the dorsal/ventral, medial/lateral arrows show in C). Striosomal D1-MSNs were omitted because their morphologies were smaller. **C)** Several of the morphological features that showed systematic variation in the coronal (samples reflected across the midline) or sagittal planes within the D1-MSN subclass (STRd D1 Matrix, STRv D1, and STRd Striosome). Asterisks indicate significance of linear regression model (* p <0.05, ** p< 0.01, *** p< 0.001), while arrow indicates direction of steepest increasing numeric value. **D)** Overlaid action potential traces for extremes of the spatial gradient (based on projection onto the axis of maximal variation in the coronal plane for upstroke) showing slower action potentials ventrally. **E)** Spatial gradients in several electrophysiological features appeared consistent across subclasses, as illustrated by the example action potential traces in panel D.

**Figure 7.**
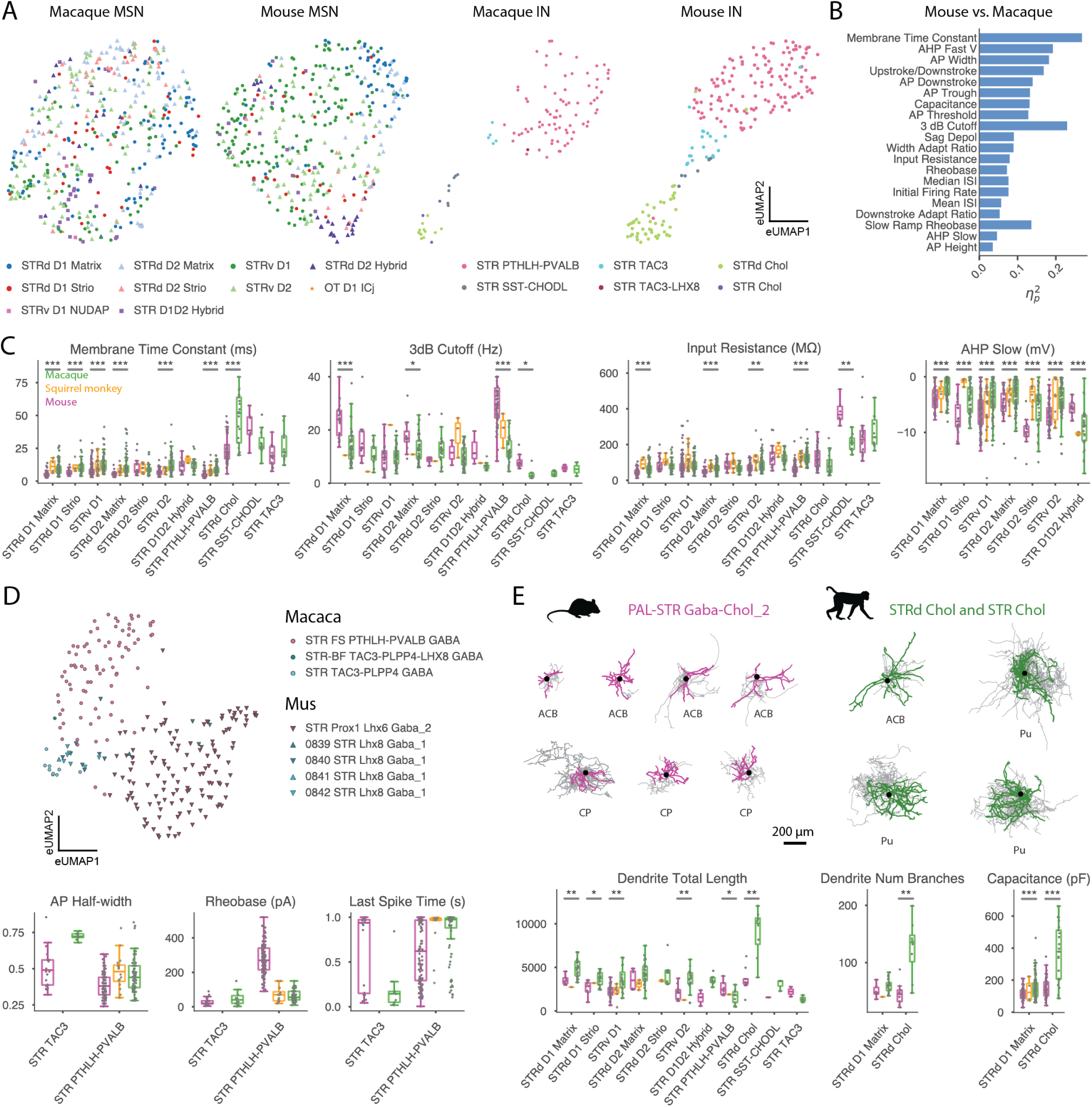
Cross-species differences in striatal cell types. **A)** UMAP visualization emphasizing common structure for MSNs and INs. Features were z-scored within each species (only mouse and macaque were included) before combining into a joint embedding. Each species is shown separately; axis ranges differ. **B)** Top species factors from two-way ANOVA comparison with species (macaque or mouse) and equivalent cell type as factors. **C)** Highlighted features that differ between mouse and macaque – statistical significance is indicated for mouse-macaque comparisons where at least three samples were available for each species. Results show slower temporal filtering, moderately increased input resistance across several groups but decreased input resistance for SST-CHODL, and a more distinctive AHP profile that sets D1D2 Hybrid neurons apart from other MSNs in primates. **D)** UMAP and feature plots highlighting differences between macaque PTHLH and TAC3 interneurons and their mouse counterparts. **E)** Example morphologies (dendrites in color, axons in gray) and quantitative analyses showing the markedly expanded and elaborated dendritic structures of Cholinergic interneurons in macaque.

We also sought to compare cortical and striatal SST-CHODL neurons, but lacked convincing samples of the rare cortical SST-CHODL type. Striatal SST neurons do appear electrophysiologically distinct from cortical SST types in general (Figure 5A), despite the relative transcriptomic proximity of cortical and striatal SST types in rodents ^23^.

#### TAC3 interneurons

The TAC3 group exemplifies pronounced transcriptomic divergence between mouse and primate striatal cell types ^26,27^. Transcriptomically, primate TAC3 interneurons correspond most closely to a subpopulation of TH+ interneurons in mouse, a small percentage of which express the *TAC3* ortholog, *Tac2*, although the primate interneurons do not express TH ^76,97^. This population represents ∼30% of striatal interneurons in primates, but despite their relative abundance, there is little known about their functionally related properties, nor how these properties compare to other better characterized striatal interneuron populations.

Morphologically, TAC3 interneurons were like PTHLH-PVALB interneurons, with both populations having similar total dendritic lengths and number of dendritic branches (Figures 4A, E and S5). Nonetheless, we identified subtle differences between the groups, including less tortuous axons in TAC3 cells (Figure S5).

In contrast, the electrophysiology of TAC3 interneurons was quite distinct from PTHLH-PVALB interneurons. TAC3 neurons exhibited spike widths intermediate between PTHLH-PVALB and SST-CHODL (Figures 4E and 5I) and the capacity to reach relatively high firing rates (Figure 5F). Their most striking characteristic was their inability to sustain firing throughout the depolarizing current step, regardless of stimulus amplitude (Figure 5F). We quantified this using the time of the last spike during the 1 s step, which robustly distinguished PTHLH-PVALB neurons from TAC3 neurons (Figure 5H). Consequently, their average firing frequency-current (f-I) relationship was much shallower than in PTHLH-PVALB neurons, despite being able to reach similar maximum instantaneous firing rates (Figure 5F, G). Therefore, PTHLH neurons had a high gain, like typical FS INs, whereas TAC3 INs had low gain and relatively rapid firing occurring only at the onset of the stimulus.

To provide insight into potential mechanisms underlying these differences, we compared the properties of single action potentials, which can reflect differences in the expression of specific ion channels ^98^. Although both cell types showed symmetrical spikes, TAC3 neurons did not exhibit the very rapid downstroke and strongly hyperpolarized fast afterhyperpolarization characteristic of PTHLH-PVALB interneurons (Figure 5I,J). These features are often attributed to Kv3 channels, ^93^ and TAC3 interneurons showed lower expression of Kv3 channel genes (*KCNC1*, *KCNC2*). TAC3 neurons also had slower upstroke kinetics, a more depolarized spike threshold and adaptation of that threshold, and the lowest amplitude action potentials of any interneuron group (Figures 5J and S6). Together, these features suggest a lower level of available inward depolarizing current. Relative to PTHLH-PVALB, TAC3 neurons expressed less of the auxiliary Na^+^ channel subunit *SCN1B* (Figure 5K), which increases peak Na^+^ current density and accelerates channel gating and recovery from inactivation ^99^. In comparison to the regular spiking SST-CHODL neurons, TAC3 neurons expressed similar levels of most Na^+^ channel alpha subunits but lacked a strong expression of *SCN9A* (Nav1.7; Figure S6). The relatively narrow spike in TAC3 neurons may in part reflect the smaller voltage excursion required to complete a shorter action potential. Genes related to K^+^ channels (*KCNA1, KCNIP4, KCNB2*) and Ca^2+^ channels (*CACNA2D3*; Figures 5K and S6) further distinguish TAC3 from PTHLH-PVALB neurons. These molecular distinctions offer potential candidate mechanisms of a moderately fast-spiking phenotype whose mechanisms differ from classical fast-spiking interneurons.

### Spatial variation in morphoelectric properties

The primate striatum contains prominent functional gradients along its anatomical axes^2,100^. Along the dorsolateral–ventromedial axis, regions shift from sensorimotor processing at the dorsolateral pole to reward and emotional regulation in the ventromedial striatum. A second gradient spans the anterior–posterior axis, transitioning from areas involved in higher-order cognitive processes anteriorly to regions specialized for motor and sensory integration posteriorly.

We assigned 3D spatial coordinates to most of our Patch-seq samples (Figure 6A), allowing us to systematically examine how morphological and intrinsic physiological properties vary within subclasses across these spatial dimensions. To quantify these variations, we fit morphoelectric properties for each subclass using linear regression models with coronal and sagittal coordinates as predictors to estimate directions of maximal variation and significance (caudate ventral tail samples were excluded due to discontinuity). For this analysis we focused on subclasses with the largest sample sizes: D1 MSN, D2 MSN and PTHLH-PVALB for electrophysiology gradients, and D1 MSN for dendritic morphology. Intriguingly, across multiple morphological and electrophysiological features, and across different subclasses, we saw a common axis of variation running dorsal, medial, and posterior from the nucleus accumbens pole. This ventromedial to dorsolateral gradient aligns with a primary gradient of gene expression seen in spatial transcriptomics ^39^.

Within the D1-MSN subclass (STRd D1 Matrix MSN, STRd D1 Striosome MSN, and STRv D1 MSN), neurons displayed maximal total dendritic length and branching dorsally, and posteriorly within the sampled extent (Figure 6B,C). Notably, input resistance increased and rheobase decreased along these axes, suggesting that the increased dendritic length in dorsal neurons contributes to their relatively low excitability (Figure S7).

Additionally, we observed systematic variation in action potential kinetics within the D1-MSN subclass that was largely mirrored within the D2 MSN subclass (STRd D2-Matrix MSN, STRd D2-Striosome MSN, and STRv D2 MSN) and STR PTHLH-PVALB neurons. In all cases the action potential upstroke was faster in dorsolateral and posterior neurons and the maximum rate of repolarization was similarly accelerated (Figure 6D, E). These differences were accompanied by narrower spike widths in dorsal neurons for D1 MSN and PTHLH-PVALB groups (Figure S7). The depth of the AHP slow component also showed spatial variation and was more pronounced for many ventral neurons, which may contribute to longer interspike intervals. In the PTHLH-PVALB subclass, ventral neurons had lower maximum firing rates (Figure S7). Although our sampling of cholinergic neurons was insufficient for formal spatial analysis, the differences we saw between the dorsal STRd and ventral STR Chol neurons were consistent with these trends. Ventral neurons had sparser morphologies and broader action potentials with slower kinetics (Figure 4H and S7). Together, these findings reveal systematic morphoelectric gradients aligned with anatomical axes, perhaps reflecting functional specialization and topographical connectivity across striatal regions.

### Cross species differences in striatal neuronal properties

Although macaque striatal neurons share broad morphoelectric similarities with rodent neurons, we directly tested whether species differences exist by comparing our macaque Patch-seq dataset to a mouse dataset collected under identical protocols (Budzillo et al., in preparation) using the homologous cell types identified in the HMBA Basal Ganglia Consensus taxonomy. We also qualitatively assessed a smaller dataset from a New World monkey (*Saimiri Sciureus*) to provide insight into variation amongst primate species. To visualize cross-species relationships in a manner that emphasizes common structure, we embedded macaque and mouse MSNs and interneurons separately, projecting each class into a shared electrophysiological feature space using UMAP, after z-scoring each electrophysiological property within species (Figure 7A). Due to limited sample size, *Saimiri* neurons were excluded from this visualization. MSNs with similar transcriptomic identities overlapped across species, but certain differences were observed, such as macaque STR D1D2 Hybrid neurons appearing more distinct than their mouse counterparts while the opposite was true for STRd D2 Hybrid neurons. However, STRd D2 Hybrid MSNs are heterogeneous, with distinct dorsal and ventral variants ^30^, and our D2 Hybrid sample was heavily biased towards dorsal regions (17/20). Interneurons with similar transcriptomic identities occupied corresponding regions of the electrophysiological embedding, exhibiting comparable relative relationships across species. This observation suggests that the general conservation of transcriptomic striatal interneuron groups in primates is reflected in broad preservation of intrinsic membrane properties ^97^.

Two-way ANOVA revealed significant species effects even after controlling for cell type (Figure 7B). Macaque neurons exhibited longer membrane time constants and, relatedly, stronger low-pass filtering (lower 3 dB cutoff), and generally higher input resistance, despite their somewhat larger dendritic trees (Figure 7B, C). *Saimiri* neurons exhibited time constants similar to macaque and longer than mouse. These properties suggest that primate neurons integrate synaptic input over longer time scales than rodent neurons. An exception to the overall input resistance trend was the SST-CHODL population: the corresponding mouse type is characterized by a very high input resistance whereas in macaque the contrast with other neuronal types is reduced.

We also observed species differences in action potential properties shared across several transcriptomic groups. Across multiple MSN and interneuron types, the spikes in macaque neurons were broader, less symmetrical, and had shorter latency and lower thresholds (Figure S8). These results are generally consistent with the findings of a recent publication comparing mouse and macaque MSNs ^12^. Finally, mouse MSNs often displayed a more pronounced slower AHP than their primate counterparts, except for the STR D1D2 hybrid (Figure 7C). The pattern of AHP slow component amplitude that distinguishes STR D1D2 Hybrid from other macaque MSNs (Figure 2) was not apparent in mouse – highlighting a primate-specific specialization.

Both TAC3 and PTHLH-PVALB interneurons exhibited striking species divergence. Unlike macaque TAC3 cells, mouse STR Lhx8 Gaba_1 neurons (the closest mouse homolog) fired continuously during current steps as quantified by their last spike times (Figure 7D), and this difference was not due to differences in the mean interspike interval at those levels of stimulation (Figure S8). Macaque TAC3 interneurons also had wider spikes and their hallmark short amplitude spikes and strong action potential height adaptation were not observed in mouse data. PTHLH-PVALB interneurons exhibited substantially lower rheobase and higher input resistance in primates. Relatedly, PTHLH-PVALB neurons were one of the few cell types that exhibited less total dendritic length and lower capacitance in macaques (Figures 7E and S8). UMAP embeddings confirmed strong separation between macaque and mouse TAC3 and PTHLH-PVALB populations (Figure 6D). Although morphological data were limited, macaque TAC3 interneurons also appeared to have more compact spherical axonal arbors as compared to more elongated and far-reaching arbors in their corresponding mouse type (Figure S8, quantified by axon max distance).

Although macaque striatal neurons generally had higher total dendritic length and capacitance (Figures 7E and S8), the difference was modest compared to differences reported for cortical pyramidal neurons ^101,102^. This contrast likely reflects anatomical constraints imposed by neocortical expansion—such as the need for dendrites to span multiple lamina—that do not apply to striatal neurons.

However, cholinergic interneurons were strikingly morphologically different across species. Macaque cholinergic interneurons exhibited a total dendritic length more than double their mouse counterparts and possessed more than twice the number of dendritic branches (Figure 7E). Although we compared our STRd Chol group to all striatal cholinergic interneurons in mouse, a similar result is seen for capacitance when limiting data to only dorsal striatal samples (Figure S8). This suggests more complex integration of their strong thalamic inputs alongside other afferent signals. In summary, species differences highlight remarkable divergence in an evolutionarily ancient structure, likely reflecting the interplay of multiple functional constraints.

## DISCUSSION

Single-cell transcriptomics has transformed our understanding of the molecular diversity of the primate striatum, revealing both cross-species similarities and differences in gene expression, spatial organization, and cell-type composition ^26,29,30,39^. However, less is known about functional properties, such as intrinsic membrane or synaptic properties, that are essential for interpreting neuronal and circuit behavior within these transcriptomic frameworks. These fundamental features are largely inaccessible from human striatal tissue, making it challenging to relate rodent studies to human health and disease. In this study, we generated a large-scale, multi-modal Patch-seq dataset from the macaque striatum. This enabled us to reveal the morphoelectric properties of living striatum neurons in a primate species, offering a translational perspective that complements the extensive rodent literature, while providing a comprehensive framework for understanding phenotypic variability. Our dataset provides validation of BICAN basal ganglia taxonomies by demonstrating strong correspondence between transcriptomic cell types and morphoelectric properties. We also found underappreciated variability in the properties of MSN and other LGE-derived neurons arising from the interacting effects of pathway, compartment, anatomical axes, and transcriptomic identity. Still, MSN properties differed across types in a relatively continuous and overlapping manner. In contrast, interneurons exhibited more discrete morphoelectric properties. Although the overall organization of striatal cell types and their relative properties resembled those in mouse, we identified several cross-species differences that may reflect primate innovations.

### Systematic, overlapping variation within MSN and MSN-like neurons

Our findings show that the continuous variability observed between transcriptomic profiles of striatal neurons is reflected in electrophysiological properties. Consistent with prior work, D1 and D2 MSNs in macaque differ only subtly in electrical properties and not necessarily in the features described for rodent. We further show that dendritic morphology is broadly similar between these canonical types. This contrasts with rodents, where morphological differences are thought to contribute to distinctions in somatic and dendritic excitability between D1 and D2 MSNs ^44^. Backpropagating action potentials decay less with distance from the soma and are more strongly modulated by dopamine in mouse D2-MSNs ^50^ – these features are thought to contribute to their increased vulnerability to spine loss following dopamine depletion. Further research is needed to understand how these less pronounced differences between primate D1 and D2 MSNs might relate to selective vulnerability in disease.

In addition to classic D1 and D2 MSNs we described the properties of several other less studied MSN and MSN-like cell types. Many distinctions between D1 and D2 matrix MSNs were also observed for the respective D1 and D2 striosome MSN counterparts, and other features differentiated striosomes from matrix MSNs in general. For example, D1-striosome MSNs exhibited a lower rheobase than their matrix counterparts, consistent with rodent data ^52^, but D2 matrix and striosome MSNs were indistinguishable in this regard. These relatively small compartmental differences suggest that the profound functional distinctions between striosome and matrix MSNs ^5,48^ arise from connectivity and neuromodulatory mechanisms rather than intrinsic excitability.

The most distinctive LGE derived neurons we characterized included non-canonical MSNs, like the D1D2 Hybrids, and ventral types like D1-ICjs. Non-canonical MSNs depart markedly from matrix and striosome MSNs in their subthreshold and suprathreshold properties, as well as ion channel related gene expression. These observations suggest that non-canonical MSNs do not correspond to exo-patch neurons – a recently discovered neuronal type in mice that has striosome like properties but are found in the matrix ^29,49^. The emerging picture is that non-canonical MSNs are not merely intermediary between different canonical MSNs but represent unique principal neuron types with distinct integrative properties and potential computational roles, a view supported by recent in vivo experiments in mice ^103^. D1 ICj granule cells, which may be involved in reward and depression ^69^, were particularly notable for their subthreshold properties and propensity for burst firing. Burst-firing is associated with enhanced capacity for driving downstream targets and engaging plasticity mechanisms ^104^ and has not previously been reported in rodent neurons ^68–70^. Because bursting can enhance the fidelity of synaptic transmission and drive large calcium influx in dendrites ^104^, these findings suggest that primate OT D1 ICj neurons faithfully communicate with their downstream targets and have enhanced windows for synaptic plasticity.

### Rich and discrete functional diversity between interneuron types

By revealing the unique morphological and electrophysiological signatures of each transcriptomic group, our findings highlight the functional modularity of striatum interneurons. In rodents, each interneuron type appears to modulate the striatal circuit in distinctive ways ^16,17^. For example, fast spiking interneurons target MSNs and other fast spiking neurons, whereas cholinergic neurons target various GABAergic interneurons ^81,105^. Subcellular targeting also varies – fast spiking neurons innervate periosomatic regions of MSNs, whereas SST-containing LTS neurons target dendrites and spines ^106^. Additionally, the unique repertoire of intrinsic conductances in each subclass endows them with distinct spontaneous and evoked firing properties and frequency preferences ^16,17,107^. Accordingly, as in rodent striatum, macaque interneurons in our dataset were highly diverse in terms of their morphoelectric properties and could be reliably distinguished based on electrophysiology alone, likely reflecting specialization for distinct circuit functions. Striatal fast-spiking interneurons differed from their cortical counterparts despite sharing some features, suggesting regional specialization of related types. Importantly though, the specific differences were distinct from those reported in mouse. Finally, the pronounced subthreshold resonance we observed in primate cortical fast-spiking neurons may reflect region-specific demands on synaptic integration and frequency tuning ^41^.

TAC3 interneurons are a divergent interneuron group sharing the same initial lineage class as TH interneurons in rodents ^76^. Despite comprising about 30% of the primate interneuron population ^26^, little is known about their functional properties. We observed that macaque TAC3 interneurons were morphologically similar to PTHLH-PVALB cells but had distinctive electrophysiological properties, including a moderately fast-spiking phenotype with short action potentials and an inability to maintain firing during sustained stimulation. These properties are reminiscent of spiking dynamics implicated in the detection of temporal synchrony of synaptic input ^108^, suggesting a potential role for TAC3 neurons in detecting changes in input correlation to the striatal circuit or for signaling transient salient events.

### Spatial variation in morphoelectric properties within discrete groups

The spatial organization of the mouse striatum has been well described, particularly along the dorsolateral–ventromedial axis, and often aligns with continuous variation in transcriptomic profiles ^28,31,109^. In our analyses, we observed a similar gradient in morphological and electrophysiological properties across features and, in some cases, concordantly for the D1 MSN subclass, D2 MSN subclass, and PTHLH-PVALB group. The gradients were not identical to those observed within rodent dorsal striatum PTHLH interneurons ^110^, and may instead reflect broader distinctions between dorsal and ventral striatum. Generally, ventral and anterior neurons had sparser morphologies and slower firing kinetics. This may reflect both a reduced degree of input integration and a slower temporal integration of limbic signals. Similarly, ventral cholinergic neurons had sparser morphologies than their dorsal counterparts, consistent with previous immunohistochemistry studies ^111^, and showed substantial electrophysiological differences, including broader action potentials and little hyperpolarization-induced sag, likely related to lower *HCN4* expression. Thus, some signature properties of dorsal cholinergic interneurons do not extend to ventral cholinergic interneurons, which are implicated in depression and its treatment ^112^.

### Cross-species divergence in properties of homologous neuronal groups

Although the striatum is an evolutionarily ancient structure, we observed substantial cross-species differences in mouse and macaque cell types, likely reflecting both adaptation and exaptation. At a coarse level, the transcriptomic alignment between mouse and macaque cell types was well supported by electrophysiological properties, yet we observed many points of divergence. Across multiple cell types, mouse striatal neurons showed less subthreshold attenuation of high-frequencies, which may better accommodate the faster contraction speeds and higher locomotor stride frequencies seen in smaller animals due to biomechanical scaling ^113,114^. Beyond the lack of conservation of many rodent D1–D2 MSN differences in macaques, STR D1D2 Hybrid neurons were particularly distinct in macaques, while a subset of D2 MSNs (corresponding to STR D2 Hybrid) appeared more distinct in mice, highlighting potential species-specialized cell types. Similarly, macaque TAC3 interneurons aligned with the closest mouse supertype (STR Lhx8 Gaba_1) but displayed notable differences from these, including distinctive phasic firing patterns and short spikes. Notably, mouse and primate PTHLH-PVALB also diverged, with macaque neurons exhibiting lower rheobase, higher input resistance, longer membrane time constant, and greater high frequency attenuation. These properties imply that primate PTHLH-PVALB neurons integrate over longer timescales, yet their lower rheobase suggests recruitment at modest input levels to normalize gain and sharpen representations. Lastly, although striatal neurons in general were only modestly larger despite the substantially larger brain size, primate cholinergic interneurons had markedly larger and more complex morphologies. Together, these distinctions provide a more robust foundation for translating rodent findings into human therapeutic contexts.

### Limitations of the study

Our sampling was insufficient to support strong conclusions about rare cell types, like D1-NUDAP and LAMP-LHX8. Likewise, limited samples sizes may have prevented us from detecting subtle differences between related groups. Our macaque cortical fast-spiking interneuron dataset was derived primarily from visual cortex and may not fully represent properties across all cortical regions.

Our sampling did not extend to the most posterior regions of the striatum, and was very limited in the ventral caudate tail, where additional variation may exist ^115^. The lack of quantitative anatomical connectivity in the anatomical reference also constitutes a crucial missing piece of context for cross-modal integration. Only local axon arborization could be considered since long range projections are truncated in the slice preparation – complementary approaches are needed to understand connectivity.

Additionally, we did not assay synaptic physiology, plasticity and neuromodulation, which are critical for circuit function. Nonetheless, the differentially expressed ion channel genes we identified provide insight into pathways engaged by neuromodulation and will support many avenues for future investigation.

Despite these limitations, this dataset—together with companion multiomic, spatial transcriptomic, viral enhancer, and other related datasets—provides a roadmap for next-generation, transcriptomics-guided studies of striatal cell types and their functional roles, helping to address these remaining gaps.

## RESOURCE AVAILABILITY

### Lead Contact

Further information and requests for resources and reagents should be directed to and will be fulfilled by the lead contact, Brian E. Kalmbach.

### Materials Availability

This study did not generate new unique reagents.

### Data and Code Availability

- Electrophysiology data in NWB format have been deposited at DANDI: https://doi.org/10.48324/dandi.001626/0.251028.1747, https://doi.org/10.48324/dandi.001465/0.250627.1703, https://doi.org/10.48324/dandi.001348/0.250401.1556, https://doi.org/10.48324/dandi.000934/0.240315.1754, https://doi.org/10.48324/dandi.000768/0.231221.2138, https://doi.org/10.48324/dandi.000635/0.230921.1734, https://doi.org/10.48324/dandi.000569/0.230706.1630
- Morphological reconstructions in SWC format have been deposited at Brain Image Library (BIL)
- Sequencing data can be found at NEMO: https://assets.nemoarchive.org/dat-nsm6mxv
- All original code has been deposited at GitHub and is publicly available as of the date of publication.
- Any additional information required to reanalyze the data reported in this paper is available from the lead contact upon request.

## ACKNOWLEDGMENTS

This publication was supported by and coordinated through the BRAIN Initiative Cell Atlas Network (BICAN) (https://braininitiative.nih.gov/research/tools-and-technologies-brain-cells-and-circuits/brain-initiative-cell-atlas-network). This work was funded by the Allen Institute for Brain Science and by the National Institutes of Health awards: UM1 MH130981 (ESL, HZ) NIH R01NS123959 (BEK, NDem), and BRAIN Armamentarium award UF1 MH128339 (JTT, TB). Its contents are solely the responsibility of the authors and do not necessarily represent the official view of the NIH, WaNPRC, or ORIP. The WaNPRC is supported by the NIH Office of Research Infrastructure Programs (ORIP) under awards P51OD010425 and U420D011123. The authors thank Madeleine Hewitt, Meghan Turner, Brian Long, Stephanie Seeman, and Jennie Close for discussions regarding spatial distribution of cell types and valuable feedback on the analyses. We thank Greg Horwitz for generously sharing his lab resources during pilot work and for detailed manuscript feedback. We thank the founder of the Allen Institute, Paul G. Allen, for his vision, encouragement and support.

## AUTHOR CONTRIBUTIONS

Tissue acquisition/processing: AJ, BEK, BRL, CE, EP, MJ, MKa, NDee, NDem, NT, TCas, XO Transcriptomic processing: CR, DB, KAS, MT, SDa, TCar

Reconstruction: ÉvT, GT, IR, JAn, KK, LA, RD, SAS

Electrophsiology: AM, BEK, GL, JTT, JTr, KBl, KH, KN, LN, MCV, MV, RR, SFO, SV, TJ Imaging/Histology: AA, AG, AH, AO, AR, CAP, JAr, JM, JW, KBe, KBr, LP, MB, MG, MMc, NID, RE, SB, SDH, STR, TE, ZCJ

Data archive / Infrastructure: ABC, ABh, BEK, BL, BRL, DR, JG, JTh, KAS, KH, MW, RD, SAS, SS, TJ, XL

Managment/Project Managment: BEK, BL, BRL, CE, ESL, GT, HZ, JTT, JW, KAS, LK, LP, NWG, SAS, SDH, TB, TJ, YF

Data analysis: ABh, ABu, BEK, BL, BRL, GT, IK, JAM, JTT, JTh, MMa, MW, NJ, NWG, RD, SFO, SS, SW, TB, TCh, TJ, XL

Data interpretation: ABu, BEK, BRL, GT, IK, JAM, JTT, NJ, NWG, RD, SLD, TB, TJ, XL, YF

Writing manuscript: BEK, JTT, RD, XL

Viral production: ACH, BEK, DN, JTT, MW, NDo, RK, SY, TB, XO Funding acquisition: BEK, ESL, HZ, JTT, NDem, TB

## DECLARATION OF GENERATIVE AI AND AI-ASSISTED TECHNOLOGIES

During the preparation of this work, the authors used OpenAI ChatGPT to debug code and ChatGPT / Microsoft Copilot to improve the readability and language of the manuscript. After using this tool, the authors reviewed and edited the content as needed and take full responsibility for the content of the publication.

## STAR Methods

### Experimental model and study participant details

#### Non-human primate specimens

We obtained non-human primate brain specimens from animals designated for the Washington National Primate Research Center’s Tissue Distribution Program. Tissue from male (n = 23) and female (n = 47) *Macaca nemestrina* (n = 59) and *Macaca mulatta* (n = 11), from 2-22 years old (9.14 ± 0.59) were used in this study. Additionally, we collected data from male (n = 2) and female (n = 3) *Saimiri sciureus* animals, aged 5-16 years old (9.75 ± 2.10). Monkeys were housed in individual cages on a 12h light/dark cycle in a temperature and humidity-controlled room. All procedures involving non-human primates were approved by the University of Washington’s Institutional Care and Use Committee (IACUC) and conformed to the NIH’s Guide for the Care and Use of Laboratory Animals.

#### Mouse specimens

Mouse samples were obtained from mixed strains of adult male and female mice. Mice had free access to food and water and were maintained on a 12-hour light/dark cycle. Mice were housed 3-5 per cage in a temperature and humidity-controlled room. All procedures involving mice were approved by Allen Institute’s IACUC and conformed to the NIH’s *Guide for the Care and Use of Laboratory Animals*. Detailed experimental procedures are described in Budzillo et al., in preparation.

### Method details

#### Brain slice preparation

We used similar brain slice preparation methods and the same solutions for all species. Under general anesthesia with isoflurane gas, non-human primate species were perfused through the heart with carbogenated (95% O_2_/5% CO_2_) artificial cerebrospinal fluid (aCSF) consisting of (in mM): 92 N-methyl-D-glucamine (NMDG), 2.5 KCl, 1.25 NaH_2_PO_4_, 30 NaHCO_3_, 20 4-(2-hydroxyethyl)-1-piperazineethanesulfonic acid (HEPES), 25 glucose, 2 thiourea, 5 Na-ascorbate, 3 Na-pyruvate, 0.5 CaCl_2_·4H_2_O and 10 MgSO_4_·7H_2_O (pH 7.4). Following exsanguination, we extracted the cerebrum from which we dissected the striatum. The striatum block was placed in chilled, continuously carbogenated NMDG aCSF and transported back to the Allen Institute for further processing. Mice were deeply anesthetized with 5% isoflurane and intracardially perfused with ∼50 mL of ice-cold aCSF, slightly different from the aCSF used for non-human primate specimens (96 NMDG, 2.5 KCl, 1.25 NaH_2_PO_4_, 25 NaHCO_3_, 20 HEPES, 25 glucose, 2 thiourea, 5 Na-ascorbate, 3 Na-pyruvate, 3 myo-inositol, 0.01 taurine, 12 N-acetyl-L-cysteine, 0.5 CaCl_2_·4H_2_O and 10 MgSO_4_·7H_2_O, pH 7.4).

We sectioned tissue blocks in cold NMDG aCSF using a vibratome (VT1200S, Leica Biosystems) or a Compresstome (VF-300, Precisionary Instruments) to produce 300-350 μm thick coronal sections of the striatum. We then transferred slices to a warmed (34°C) and carbogenated chamber containing NMDG aCSF for 10 minutes after which slices were transferred to a chamber containing room temperature aCSF consisting of (in mM): 92 NaCl, 2.5 KCl,1.2 NaH_2_PO_4_, 30 NaHCO_3_, 20 HEPES, 25 D-glucose, 2 thiourea, 5 sodium-L-ascorbate, 3 sodium pyruvate, 2 CaCl_2_·4H_2_O and 2 MgSO_4_·7H_2_O. Slices destined for acute brain slice recordings were maintained in this aCSF at room temperature until the time of Patch-seq experiments.

We reserved a subset of non-human primate brain slices from each tissue block for organotypic slice culture. We placed slices onto membrane inserts (Millipore) in 6 well plates containing 1 mL per well of culture medium (8.4 g/L MEM Eagle medium, 20% heat-inactivated horse serum, 30 mM HEPES, 13 mM D-glucose, 15 mM NaHCO_3_, 1mM ascorbic acid, 2mM MgSO_4_·7H_2_O, 1 mM CaCl_2_·4H_2_O, 0.5 mM GlutaMAX-I and 1 mg/L insulin). We adjusted pH and osmolality of the slice culture medium to pH 7.2-7.3 and 300-310 mOsmoles/kg by adding sterile, filtered H_2_O. Stock culture medium was stored at 4°C for up to 2 weeks. Culture plates were stored in a humidified 35°C, 5% CO_2_ incubator and culture medium was replaced every 2-3 days. In a subset of cultures, we directly applied concentrated adeno-associated virus (AAV) vectors 1-12 hours after plating to express a fluorophore in the cell types of interest. The use of slice cultures also extended the life of precious tissue specimens several days beyond the viability of acute brain slices.

#### Patch-seq recordings

A detailed protocol for the Patch-seq methods we used in this study can be found at protocols.io (https://dx.doi.org/10.17504/protocols.io.bw6gphbw). Before data collection, surfaces and equipment were thoroughly cleaned with DNA away (Thermo Scientific), followed by RNAse Zap (Sigma-Aldrich) and nuclease free water. Brain slices were placed in a submersion recording chamber and continuously perfused with fresh, temperature controlled (32-34°C) recording aCSF consisting of (in mM): 126 NaCl, 2.5 KCl, 1.25 NaH_2_PO_4_, 26 NaHCO_3_, 12.5 glucose, 2 CaCl_2_·4H_2_O and 2 MgSO_4_·7H_2_O, 1 kynurenic acid and 0.1 picrotoxin (pH 7.3-7.4). We visualized neurons with upright microscopes (Olympus BX51WI or Scientifica SliceScope Pro) equipped with 40x water immersion objectives and infrared differential interference contrast optics or oblique illumination optics.

We pulled patch clamp pipettes from thick-walled borosilicate glass (Warner Instruments, G150F-3 or King Precision Glass type 8250) using an electrode puller (Narishige PC-10 or Sutter Instruments P-87). The open tip resistance was 4-6 MΩ when filled with 2 µL of the internal recording solution consisting of (in mM): 110.0 K-gluconate, 10.0 HEPES, 0.2 EGTA, 4 KCl, 0.3 Na_2_-GTP, 10 phosphocreatine disodium salt hydrate, 1 Mg-ATP, 20 mg/mL glycogen, 0.5U/mL RNase inhibitor (Takara, 2313A), 0.5% biocytin and 0.02 Alexa 594 or 488 – pH adjusted to 7.3 with KOH. We acquired whole cell somatic recordings using a Multiclamp 700B (Molecular Devices) amplifier and custom acquisition software (MIES, https://github.com/alleninstitute/mies) written in Igor Pro (Wavemetrics, RRID:SCR_000325). Electrical signals were digitized at 50 kHz by an ITC-18 (HEKA) and digitally low-pass filtered at 10 kHz (Bessel). Data are reported uncorrected for the measured –14 mV liquid-junction potential.

Following the acquisition of electrophysiology data, we centered the pipette towards the nucleus and applied negative pressure (–30 – –90 mbar) to extract cytosol and coax the nucleus into the pipette opening. While maintaining negative pressure we then retracted the pipette diagonally in the x,z direction, while monitoring the seal resistance and confirming that the nucleus was following the path of the pipette. We then expelled the nucleus, cytosol and internal solution into a PCR tube containing lysis buffer (Takara, 634894) for further processing.

#### Processing of Patch-seq samples

We used SMART-Seq v4 Ultra Low Input RNA kit for Sequencing (Takara, 634894) to reverse transcribe poly(A) RNA and amplify full-length cDNA for 20 PCR cycles according to manufacturer’s instructions. Samples proceeded through library construction using Nextera XT DNA Library Preparation Kit (Illumina FC-131-1096). Nextera XT DNA Library prep was performed according to manufacturer instructions except that the volumes of reagents including cDNA input were decreased to 0.2x by volume. Samples were sequenced to approximately 0.5-1 million paired-end 50b reads/sample.

Paired end reads from macaque samples were aligned to the Mmul_10 rhesus macaque reference genome (https://www.ncbi.nlm.nih.gov/datasets/genome/GCF_003339765.1/) using a RefSeq annotation gff file retrieved from NCBI (annotation release 103). Squirrel monkey samples were aligned to BCM_Sbol.2.1 reference genome (GCF_016699345.2) using a RefSeq annotation gff file retrieved from NCBI (annotation release 102). We performed sequence alignment using STAR (v2.5.3, RRID:SCR_004463) in two-pass mode. Only uniquely aligned reads were used in gene quantification. We quantified exonic and intronic reads separately using the R Genomic Alignments package summarizeOverlaps function using Intersection-NotEmpty mode.

#### Biocytin histology

Following Patch-seq experiments, we fixed brain slices in 4% paraformaldehyde in 1x phosphate buffered saline (PBS) for 1-2 days followed by PBS at 4⁰ C until staining. We used a horseradish peroxidase enzyme reaction using diaminobenzidine (DAB) as the chromogen to visualize biocytin-filled neurons. Slices were also stained with DAPI and incubated in 1% hydrogen peroxide for 30 minutes. Following permeabilization with 2% Triton-X in PBS, we incubated slices in ABS with 0.1% Triton at 4⁰ C overnight or up to two days. We then rinsed slices in PBS three times.

Slice overview images of mounted sections were acquired on either an upright AxioImager Z2 microscopes (Zeiss, Germany) equipped with an Axiocam 506 camera and a 20x objective (Zeiss Plan-NEOFLUAR 20x/0.5), or on a SLIDEVIEW VS200 (Evident Scientific, Japan) equipped with a Hamamatsu ORCA-Fusion-BT camera and 10x objective (Evident UPLXAPO10X). Single-plane DAPI images and brightfield transmission z-stacks (36–64 μm total depth, 8–12 μm spacing) were collected, and z-stacks were collapsed into minimum-intensity projections.

After cells were selected from slice overview images, tiled z-stacks of individual cells were acquired in brightfield for the purpose of automated and manual reconstruction using Axioimager Z2 microscopes equipped with 63x objectives (Zeiss LD LCI Plan-Apochromat 63x/1.2 Imm Corr or Zeiss Plan-Apochromat 63x/1.4 Oil) at z-step size of 0.44 μm. Tiled images were stitched in Zeiss’ ZEN software and exported as individual-plane TIFF files.

#### Morphological reconstructions

For a subset of neurons with good quality transcriptomics we generated reconstructions of dendrites and in some cases axons from the 63x image stacks. Image stacks proceeded through our Vaa3D-based image processing and reconstruction pipeline (Ref). We used TreMAP (Ref) to generate an automated reconstruction of each neuron, which was then manually proofread in the Mozak extension in Vaa3D.

#### Anatomical annotations

We pinned Patch-seq samples to a macaque reference brain volume (Mac25Rhesus_v2) annotated with the Harmonized Ontology of Mammalian Brain Anatomy (HOMBA) basal ganglia segmentation. Block images obtained during dissection, individual slice images obtained during brain slice preparation and slice overview images obtained after biocytin staining were used to place each slice in 3D space by matching photo-documentation with the segmented reference in slicer (https://www.slicer.org/). We then pinned individual cells within this reference, while noting the anatomical nomenclature and the x, y, z coordinates within the segmented reference.

Available dissection, block face, and slice overview images were used to place macaque tissue slices in approximate 3D space by aligning block-level photodocumentation to a macaque rhesus MRI reference volume “Mac25Rhesus_v2_AverageT1w_restore_0.16mm.nn.gz”, using the 3D Slicer interactive platform (*). The “Mac25Rhesus_v2_AverageT1w_restore_0.16mm” volume and “D99Atlas_v2_Subcortical_RegByRIKEN1_0.16mm” segmentation were loaded alongside corresponding structure label maps “D99_labels_3dslicer.txt” to enable 3D anatomical visualization.

For each tissue block, images from the donor specimen were examined to determine anterior-posterior boundaries. These boundaries were matched to the MRI volume by scrolling through the coronal axis (Y-dimension) in 3D Slicer to identify corresponding y-coordinates for each bookend slice. Once anterior-posterior block boundaries were determined, interleaved sections were assigned a y voxel (1 voxel = 160 um) based on anatomical landmarks, allowing for approximate registration of each physical slice within the MRI volume. Dissection angles were estimated by inspecting photodocumentation for deviations from coronal slicing and corrected within the “Reformat” module of 3D Slicer to match off-coronal slice angles.

Once slice positions were established, individual cells were annotated using the Markups module in 3D Slicer. A new Point List was created for each slice and populated with control points corresponding to the location of each individual neuron, as determined from the annotated slice overview image. The hemisphere was determined using available dissection images and if it could not be determined right hemisphere was assigned as default.

#### Morphological reconstruction

Reconstructions were generated based on 63X image stacks described above. Stacks were run through a Vaa3D-based image processing and reconstruction pipeline. An automated reconstruction of some neurons was generated using the approach described in Gliko 2022. Automated or manually initiated reconstructions were then extensively manually corrected and curated using a range of tools (for example, virtual finger and polyline) in the Mozak extension (Zoran Popovic, Center for Game Science, University of Washington) of Terafly tools in Vaa3D.

#### Adeno-associated virus (AAV) vectors, cloning and packaging

We used an AAV vector (AiP13445, pAAV-AiE0873m-minBG-SYFP2-P2A-3XFLAG-10aa-H2B-WPRE3-BGHpA) previously demonstrated to label cholinergic interneurons in the striatum of mice and macaques following *in vivo* application (Hunker et al., 2025). Full plasmid sequences and maps are available at Addgene (plasmid ID: 191725). Additionally, we used two vectors described in our companion manuscript ^116^, AiP14609 (pAAV-AiE1223q-minBG-SYFP2-WPRE3-BGHpA) and AiP15223 (pAAV-AiE1476q-minBG-SYFP2-WPRE3-BGHpA), to target granule cells in the islands of Calleja.

We packaged PHP.eB serotyped AAV vectors following a previously established protocol ^117^. Crude AAV vector was used for direct application to the surface of macaque striatum slices cultured on membrane inserts as described above.

### Quantification and statistical analysis

#### Neurophysiology features

The analysis we present are based on 716 macaque samples and 96 squirrel monkey samples that passed transcriptomic QC. Of these, 33 macaque samples were from a previously published study^12^ that included RNA-sequencing but no formal mapping to a reference taxonomy. Additionally, we included 667 mapped samples from mice from another study (Budzillo et al., in preparation). For comparison to cortical neurons, we included macaque cortical samples totaling 63 Pvalb and 7 Sst post-QC samples. We measured physiological features from voltage responses elicited by four main stimulus sets: 1) a 3 ms suprathreshold current pulse 2) a series of 1 s long suprathreshold current steps 3) a series of 1 s long hyperpolarizing current steps and 4) a chirp stimulus. For suprathreshold current injections we used online analysis functions to measure rheobase within 10 pA. For the 1 s current injections we also probed the spiking behavior of neurons in response to current injection amplitudes 30-130 pA beyond rheobase in 20 pA current steps. In some neurons we used online analysis functions to dynamically adjust this series of current steps to probe spiking up to current amplitudes that produced Na^+^ channel inactivation-mediated spike failure. The hyperpolarizing current injections consisted of a series of 1 s square wave steps that varied from –30 to –110 pA in 20 pA steps. For neurons with very high input resistance (e.g. OT D1-ICJ granule cells), we scaled these current injections down. The chirp stimulus consisted of a constant amplitude sinusoidal current injection that increased in frequency logarithmically from 0.2-40 Hz in 20 s. The amplitude of the chirp was scaled to produce a peak-to-peak maximum voltage deflection of 6-10 mV.

We measured subthreshold and suprathreshold properties as previously described ^33^. Briefly, we detected action potential thresholds elicited during the 1 s and 3 ms long steps by identifying time points where the smoothed derivative of the membrane potential exceeded 20 mV/ms. We then measured each action potential’s height (threshold-to-peak), width (at half-height), after-hyperpolarization (relative to threshold), maximum rates of depolarization (upstroke) and repolarization (downstroke) and the ratio of the upstroke and downstroke. From these suprathreshold stimulus protocols we also calculated rheobase, slope of the firing rate versus current amplitude relationship, the first spike latency relative to the onset of the current step and initial firing rate (1/the first inter-spike interval) at rheobase, and the mean ratio of consecutive inter-spike intervals at ∼50 pA above rheobase. From the hyperpolarizing current steps we measured subthreshold features including input resistance, membrane time constant and sag fraction which was calculated as the difference in the membrane potential between the minimum value and the steady-state value divided by the peak deflection during the stimulus, ranging from 0 (no sag) to 1 (complete return to the resting potential). Adapt ratios were calculated as the feature value for the fourth spike relative to first spike. We constructed impedance amplitude profiles (ZAP) from the ratio of the Fourier transform of the voltage response to the Fourier transform of the chirp 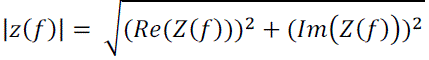. where Im(Z(f)) and Re(Z(f)) are the imaginary and real parts of the impedance Z(f), respectively. We defined resonance frequency as the frequency at which the maximum impedance occurred and resonance strength as the ratio of the maximum impedance to the impedance amplitude at .2 Hz. The 3dB cutoff was defined as the frequency at which the ZAP attenuated to a value of 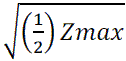. We also generated impedance phase profiles (ZPP) as 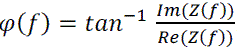. Synchrony phase was the frequency at with the ZPP was 0. We measured resting membrane potential upon obtaining whole cell configuration. Features were calculated using the IPFX python package (https://github.com/AllenInstitute/ipfx).

We applied quality control criteria for each sweep as previously described ^38^ using online analysis functions. In brief, sweeps were included for analysis if: 1) the membrane potential at the start of the sweep was within 2 mV of the initial resting potential, 2) the current required to keep the membrane potential at the original membrane potential was ± 100 pA and, 3) the root mean square noise was less than 0.2 mV for high frequency noise and 0.5 mV for low frequency noise. Sweeps failing these criteria were terminated and the same stimulus was applied again until satisfying these criteria. The intrinsic properties of some cell types made it difficult to pass online, sweep level QC. Cell types that fire action potentials spontaneously, like Chat cells (Refs), were especially difficult in this regard. In these cases, we applied a negative bias current to hyperpolarize the cell to ∼-65 mV to quiet spiking, ran similar stimulus sets without online QC and assessed sweep quality offline. Finally, for each recording we included cells in analysis if the seal before break in was greater than 1 GΩ and if the initial access resistance was less than 20 MΩ or less than 25% of the input resistance. We manually examined cells that failed these criteria for recording quality and manually passed or failed them.

OT D1-ICj granule cells were very small and challenging to patch, extract the nucleus, and maintain the intactness for morphological recovery. Often, the experimentalists used small-tipped pipettes that required a relaxation of the 20 MΩ maximum allowed access resistance for the Patch-seq pipeline in general. The actual mean access resistance for our OT D1 ICj recordings was 26.2 ± 2.7 MΩ.

#### Patch-seq mapping

We mapped non-human primate Patch-seq samples to the HMBA Basal Ganglia Consensus macaque taxonomy ^30^ and mouse Patch-seq samples to a recently generated mouse whole-brain taxonomy ^32^. To identify the cell type of the Patch-seq sample we used the Hierarchical Approximate Nearest Neighbor (HANN) method implemented in the Allen Institute MapMyCells python package (https://github.com/AllenInstitute/cell_type_mapper and https://brain-map.org/bkp/analyze/mapmycells). At each node of the taxonomy HANN mapping selects marker genes that differentiate each child of the node and then finds the approximate nearest neighbor of the Patch-seq sample using marker gene correlation as the distance metric. Most data analysis was performed at the “Group” level of the taxonomy which is the lowest level at which clear cross-species correspondences were established.

Macaque cortical samples were mapped to the “Great Apes” macaque cortical taxonomy ^118^.

#### Mapping QC

We applied additional transcriptomic inclusion criteria based on both sample-quality and mapping-quality metrics. Samples had to satisfy 1) percent reads aligned ≥ 25 and 2) genes detected ≥ 1000. For macaque samples, we additionally enforced the following criteria on mapping metrics: “Group correlation coefficient” ≥ 0.32 and “Group aggregate probability” ≥ 0.4. Because the optimal cutoff appeared different across cell types, we selected thresholds that balanced excluding spurious mappings with retaining reasonably mapped cells. For saimiri samples, which were mapped to the macaque taxonomy, we relaxed the mapping metric criteria to: “Group_correlation_coefficient” ≥ 0.15. For cortical samples, we applied the mapping criteria “Normalized Marker Sum” ≥ 0.55. Detailed quality control procedures for the mouse data are described in Budzillo et al., in preparation.

#### Morphology feature analysis

Prior to morphological feature analysis, reconstructed neuronal morphologies were expanded in the dimension perpendicular to the cut surface to correct for shrinkage after tissue processing. The amount of shrinkage was calculated by comparing the distance of the soma to the cut surface during recording and after fixation and reconstruction. For primate data, in cases where this shrinkage expansion value was in the 99th percentile of human slices, the average human shrinkage correction value (2.088) was used. Features predominantly determined by differences in the z-dimension were not analyzed to minimize technical artifacts due to z-compression of the slice after processing.

We calculated features using the skeleton keys (https://github.com/AllenInstitute/skeleton_keys) and neuron morphology (https://github.com/AllenInstitute/neuron_morphology) python packages.

#### Data analysis

Data were analyzed using custom scripts in R and Python. Since not all features were available for each cell, the data were imputed using k-Nearest Neighbors for features and samples that were at least 80% data complete prior to analyses that did not tolerate null values such as UMAP embedding or classification.

#### Classification analysis

Classification was performed using one-versus-rest logistic regression implemented in Python’s scikit-learn package in “balanced” mode to prevent overrepresented classes from dominating the model. Several other machine learning models were tested but did not show substantially better performance, so logistic regression was selected for its interpretability. Performance was then assessed via leave-one-out crossvalidation.

#### Gene expression analysis

A list of 162 ion channel-related genes was collated from the HUGO Gene Nomenclature Committee’s gene group reports (e.g., “<u>Sodium voltage-gated channel alpha subunits</u>”) to assess their expression profiles for cell-type-specific patterns. The FindMarkers() workflow in Seurat was used for discovery of differentially expressed genes between cell populations. Significance plotted on figures is based on Welch’s t-test on the counts per million values for all pairs of groups shown, with Benjamini–Hochberg correction for the number of pairs.

#### Statistical analysis

We applied outlier removal to the electrophysiological features. First, we looked at the distributions for features that should logically be positive or negative (e.g., time intervals and time constants, slope of action potential depolarization or repolarization) and removed the few points that had the wrong sign. If a feature’s distribution had skewness greater than 4, a log transformed version was created to improve normality. For each combination of feature (log transformed if necessary) and group, data points that were greater than 2.5 times the interquartile range above Q3 or below Q1 were considered extreme outliers and removed. If more than 2% of the values for a given feature were detected as outliers, the threshold was increased incrementally until it fell below 2%, preventing excessive trimming of atypical distributions.

To test for statistical significance we first performed one– or two-way ANOVA to identify features that showed significant group differences. Features were transformed if needed and the Benjamini-Hochberg false discovery rate (BH-FDR) was controlled at alpha = 0.05). Only features with at least two samples per unique group (e.g., species × cell type combinations for Figure 7) were included; if necessary, one cell type with insufficient samples was excluded to meet this criterion, except in the case of the D1/D2 matrix/striosome comparison. When displaying ANOVA results in figures, the same feature computed on various types of sweeps (e.g., short or long square steps) were consolidated for brevity. Post-hoc comparisons were conducted using Welch’s t-tests for all pairs of groups with at least 3 samples, again applying BH-FDR correction. Resulting significant pairs (p < 0.05) are indicated above the boxplots. For cross-species comparisons, only mouse and macaque comparisons are shown.

#### Cytosplore Viewer for integrated multi-modal visual patch-seq exploration

To visualize the multi-modal patch-seq data in an integrated manner, we developed a project file for the Cytosplore Viewer visualization software (available at https://viewer.cytosplore.org). This project file is tailored to the specific analysis and integration questions addressed with the basal ganglia patch-seq data in this paper, and includes the data. Cross-modal exploration and analysis of the electrophysiological and morphological features and of embeddings is realized through multiple linked visualizations. A screenshot and description of linked views is provided in supplementary figure S1. Cell groups of interest can be selected either through associated metadata (e.g., by subclass or supertype) or via free-form brushing within the morphology and electrophysiology embeddings. Selected identical cells are simultaneously highlighted in all views. For cells with available morphological reconstructions, three-dimensional morphologies are displayed alongside a representative electrophysiological voltage trace. Additional visualizations support numerical feature analysis for both the morphology and electrophysiology data, highlighting which features distinguish the selected cells from the rest of the data. This enables straightforward visual comparison between the morphologies of the cells and their electrophysiological properties. Together, the Cytosplore Viewer project file integrates bi-modal comparison of cell groups and facilitate the identification of within– and across-type differences through both visual inspection and quantitative feature analysis. This provides an integrated view on the cell-type specific characteristics.

---

**Figure S1.**
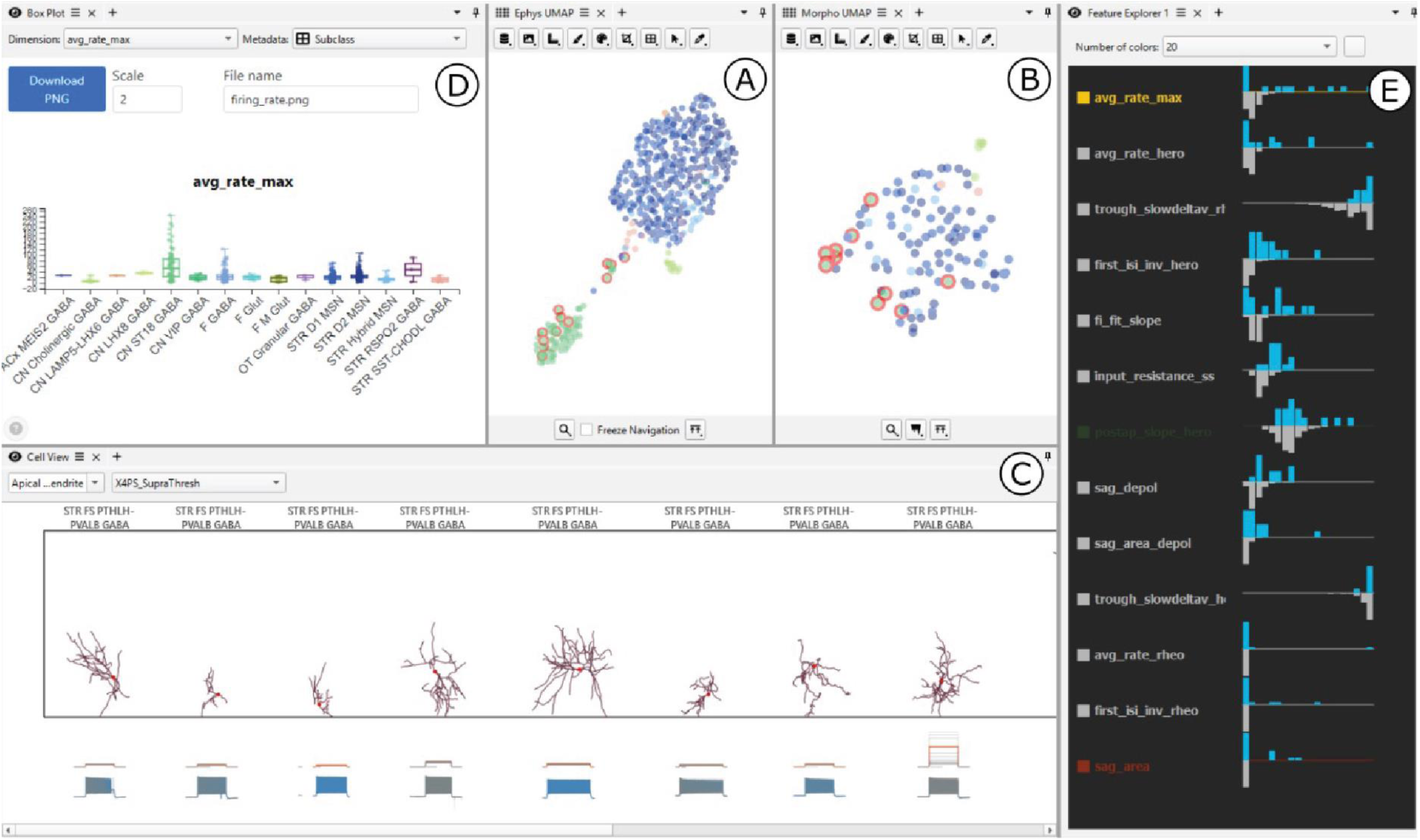
Cytosplore viewer enables for the exploration of morphoelectric properties of transcriptomic cell types in the macaque striatum, related to Figure 1. **A)** UMAP of electrophysiological features for the subset of cells with available electrophysiological recordings. **B)** UMAP of morphological features for the subset of cells with available morphological reconstructions. **C)** Visualization of the morphologies and a corresponding representative voltage trace of the selected cells, when available. Different stimulus protocols can be selected to display their associated voltage responses. **D)** Box-and-whisker plot of individually selectable data features, overlaid by the individual data points. Data points (cells) are grouped and color-coded according to their group level assignment in the taxonomy. **E)** Ranked list of either morphological or electrophysiological features, ordered from most to least salient for the selected cells. For each feature, the blue histogram shows the distribution of values for the selected cells, while the grey histogram shows the distribution across the entire dataset. This example highlights that the selected cells exhibit a particularly high firing rate relative to other cells. This is corroborated by view **C)** and **D)**.

**Figure S2.**
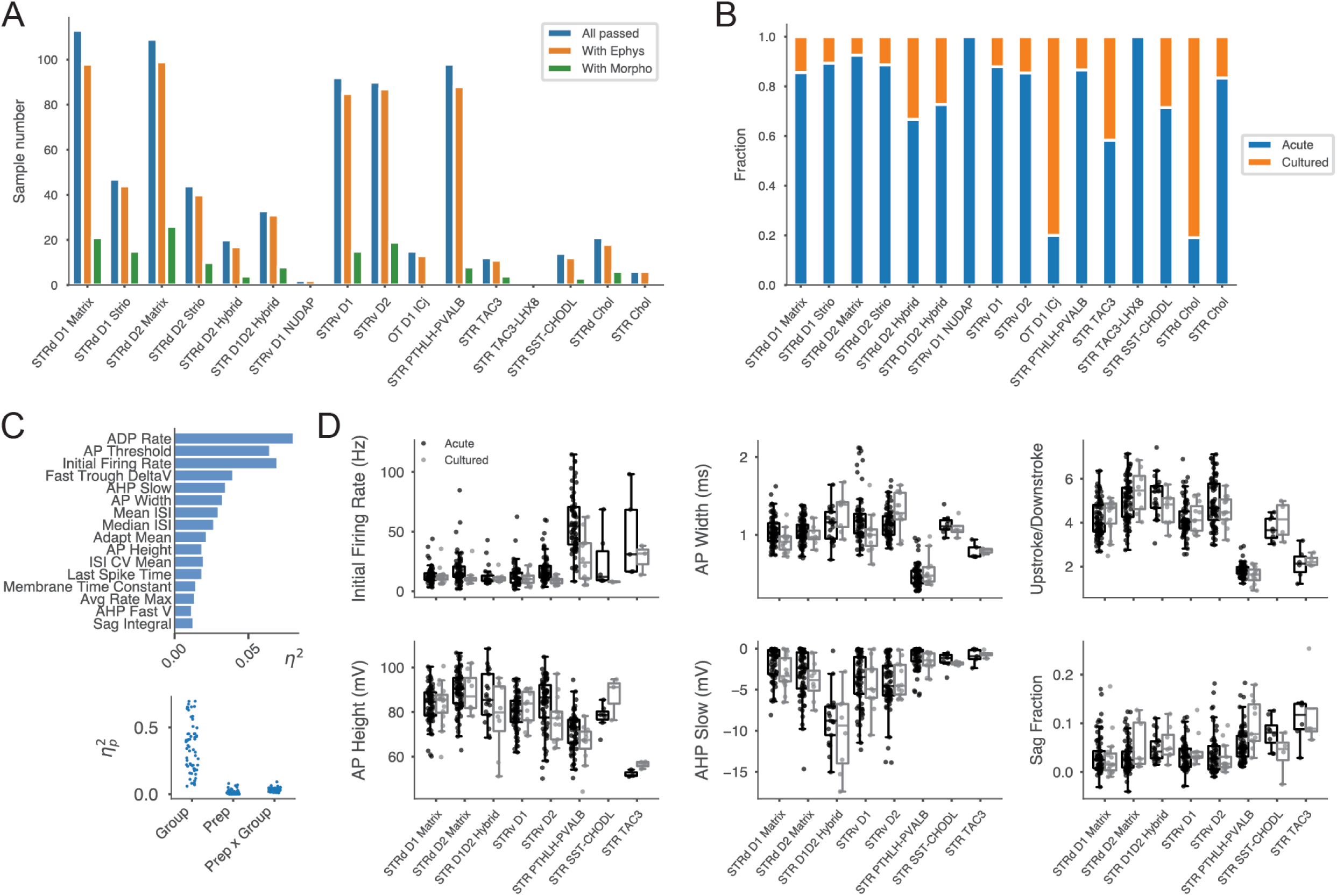
Additional dataset information, related to Figures 1 and 2. **A)** Sample sizes by group, showing total passing cells and the subsets with electrophysiological or morphological features. **B)** Breakdown of samples by preparation type (acute vs culture). **C)** Effect sizes for features differential between acute and culture samples in a two-way ANOVA with transcriptomic Group and Preparation as factors (top) for all neuron types with sufficient samples in each preparation type. Partial eta squared values for Group and Preparation, as well as their interaction; difference between transcriptomic Groups were generally much larger than the effects of culture (bottom). **D)** Example features that appear in the results – some of which show culture-dependent effects (panel C) – partitioned by culture versus acute.

**Figure S3.**
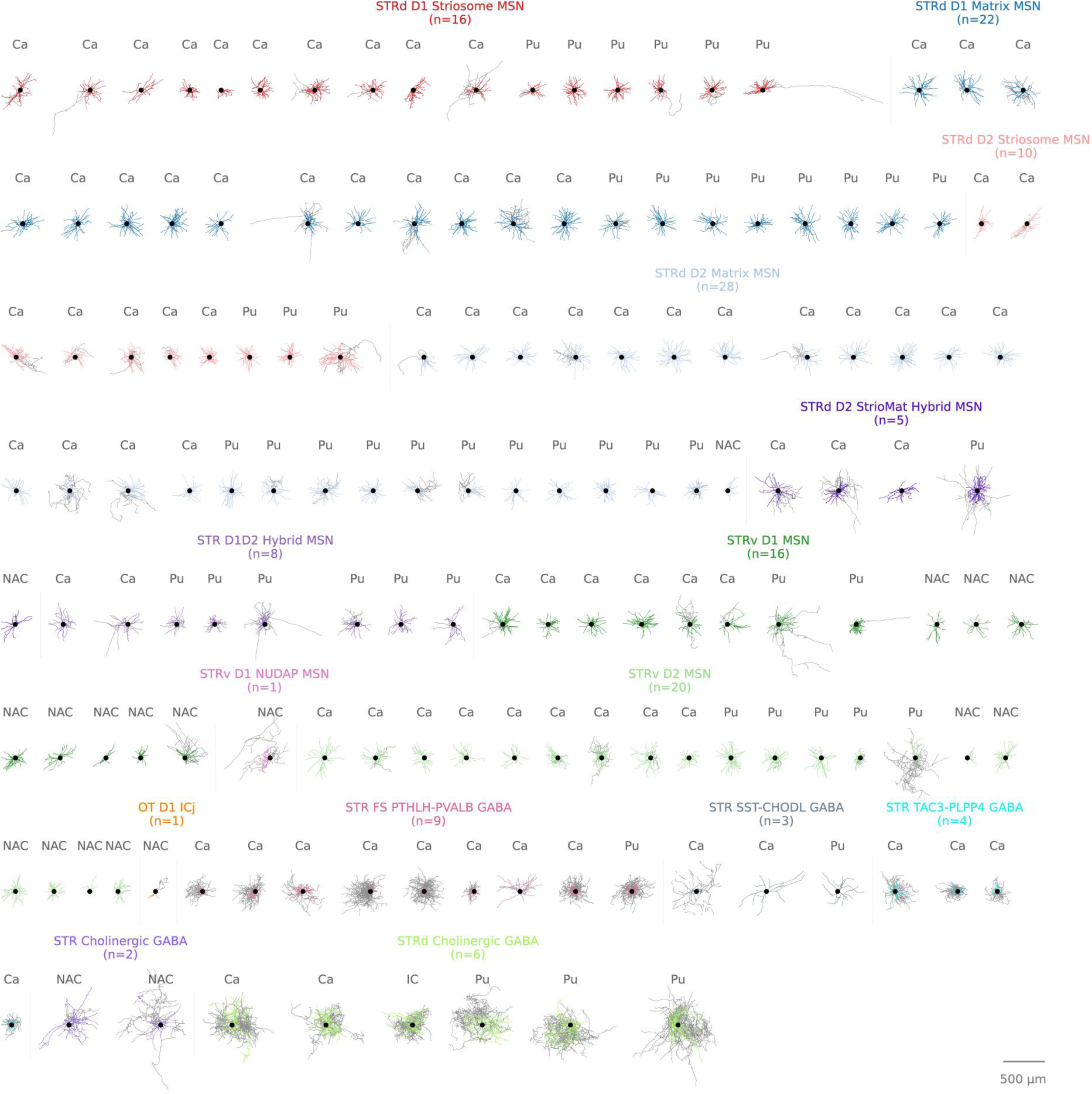
Morphologies of all reconstructed neurons, related to all figures. Dendritic processes are color coded by transcriptomic group and axons are gray.

**Figure S4.**
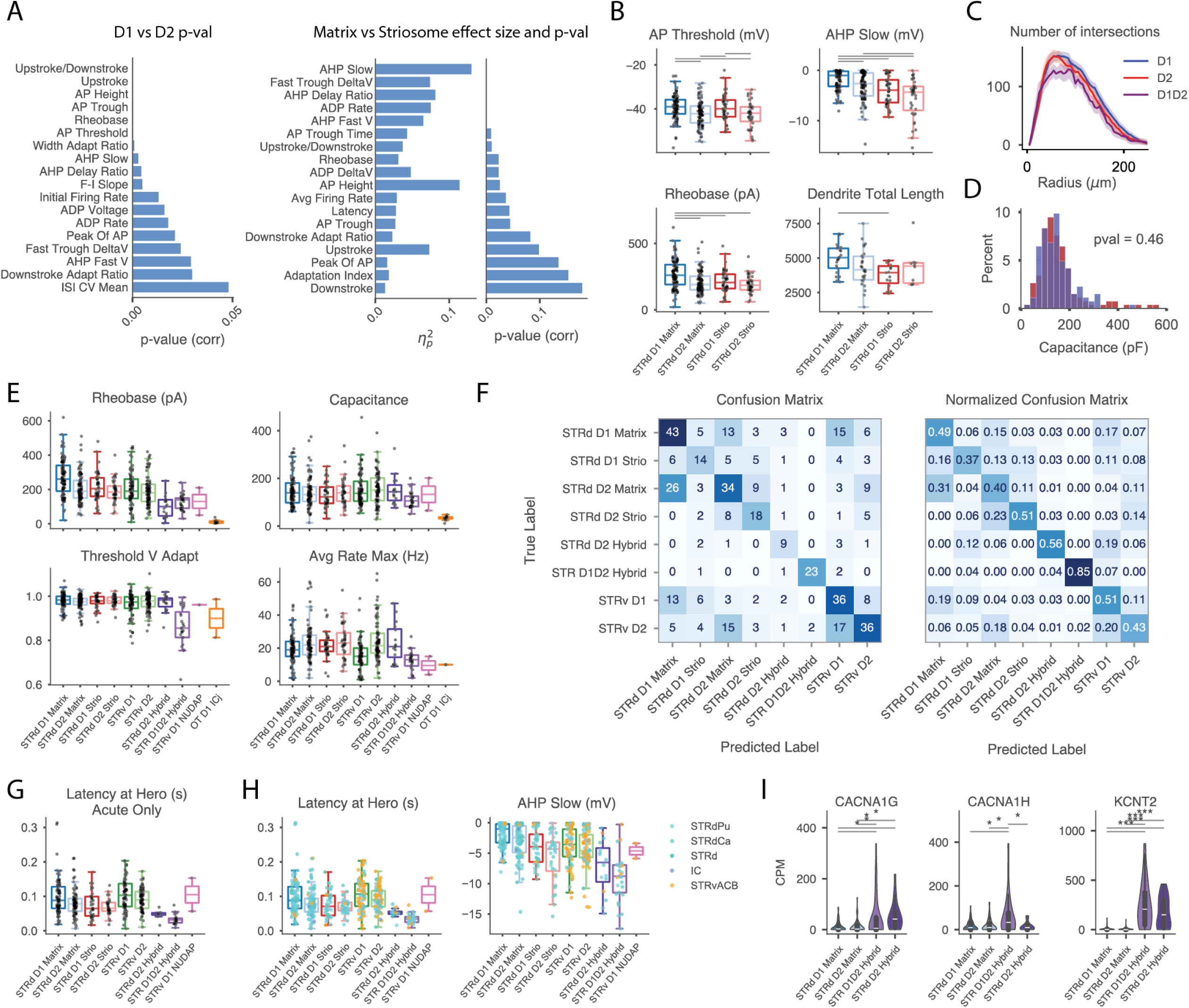
Additional morpho-electric comparisons for MSN and MSN-like neurons, related to Figures 2 and 3. **A)** For the two-way ANOVA of D1 vs. D2 and matrix vs. striosome shown in Fig. 2D, *p*-values for D1 vs. D2 and effect sizes and *p*-values for matrix vs striosome. **B)** More properties that differed on average between D1 and D2 MSNs and/or matrix and striosome. **C)** Sholl plot showing largely similar morphological properties for D1 (n = 33) and D2 MSNs (n = 25) and somewhat more compact arbors for STR D1D2-Hybrid MSNs (n = 7). Matrix and striosome cells were combined. **D)** Histogram of calculated capacitance values for D1 and D2 MSNs; the independent t-test did not reveal a significant difference between the group means. **E)** Additional features that were contrastive between main MSNs and non-canonical MSNs. **F)** Test classification performance for leave-one-out validation in a logistic regression classifier trained to categorize MSN neurons. **G)** Spike latency ∼40 pA above rheobase (“hero”) for acute samples only – a similar contrast is seen for STR D1D2 Hybrid MSNs as when considering both acute and cultured samples. **H)** Two features that distinguish STR D1D2 Hybrid MSNs from canonical MSNs, with sample region indicated by color; regional origin does not account for the effects. **I)** Additional differences in ion channel gene expression between canonical and non-canonical MSNs.

**Figure S5.**
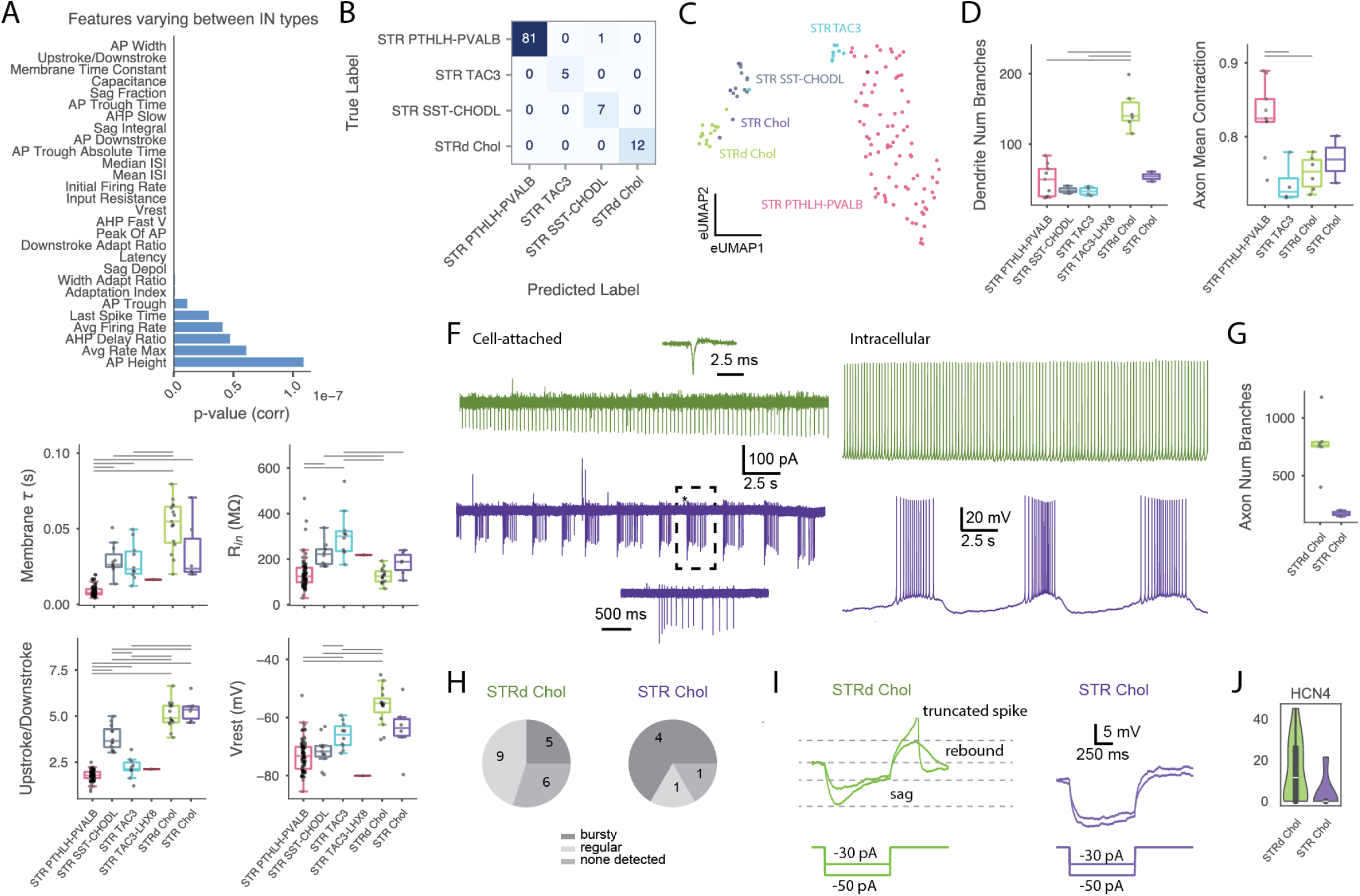
Extended figures pertaining to interneurons in general and cholinergic interneurons in particular, related to Figure 4. **A)** p-values for ANOVA shown in Fig. 4D. **B)** Unnormalized counts for classifier confusion matrix shown in Fig. 4C. **C)** UMAP of electrophysiological features generated using only striatal interneuron types (cortical neurons excluded). The embeddings in the main figures show only striatal neurons but were computed including some cortical interneuron types. **D)** Additional dendritic and axonal features distinguishing interneuron types. **E)** Additional electrophysiological features that differentiate interneuron types. **F)** Cell-attached voltage clamp and intracellular current clamp recordings of spontaneous activity in STRd Chol (dark green) and STR Chol (purple) interneurons, showcasing both regular and bursting firing patterns. **G)** Number of axonal branches in STR Chol and STRd Chol neurons. **H)** Prevalence of spontaneous activity, either bursting or regular, in cholinergic interneurons. **I)** Representative traces of hyperpolarizing sag (or lack thereof) in STRd Chol and STR Chol neurons. **J)** Expression of *HCN4*, a subunit contributing to I_h_ current with slow kinetics, is markedly lower in STR Chol (units in counts per million).

**Figure S6.**
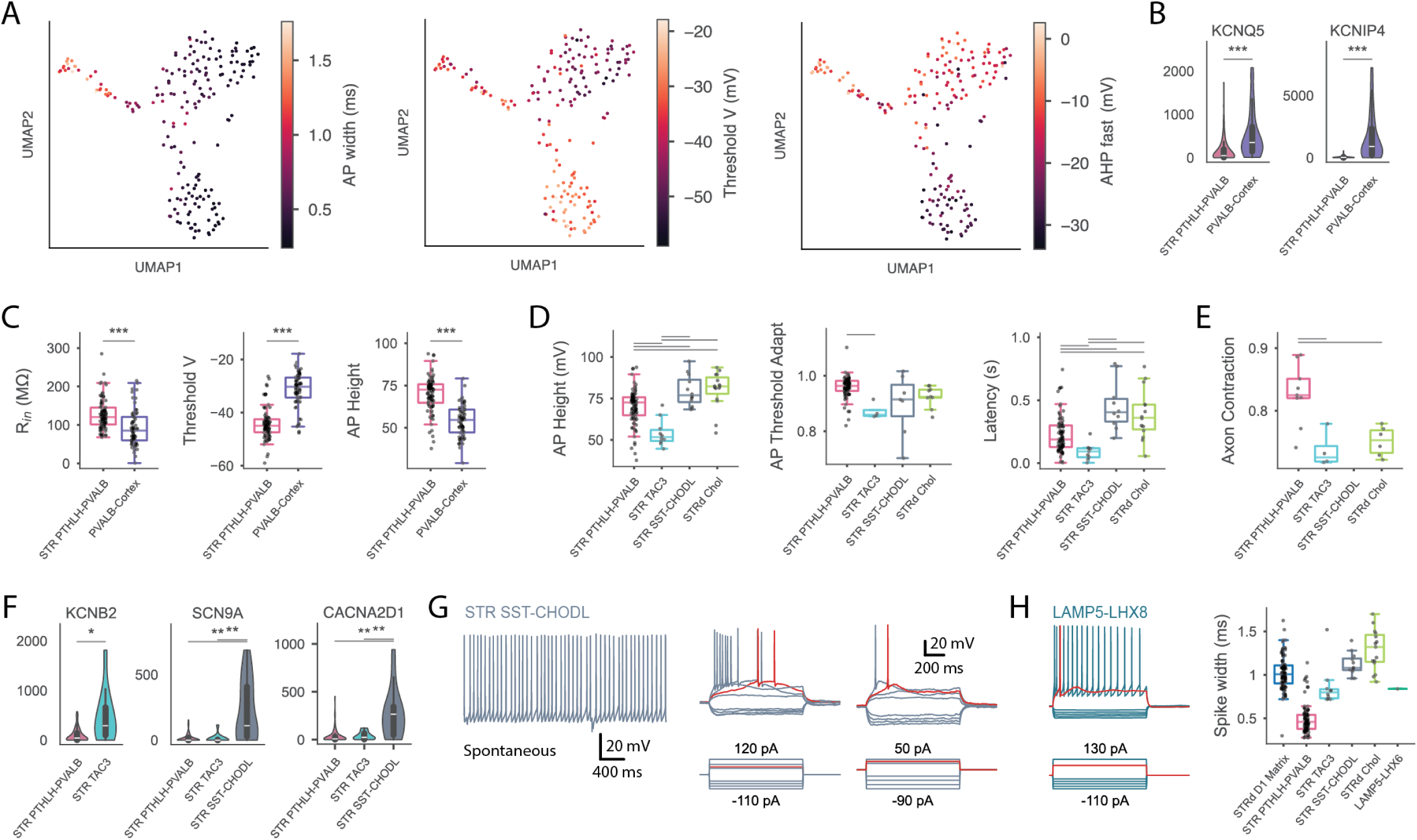
Further details on PTHLH, cortical FS interneurons, STR SST-CHODL, and LAMP5-LHX8, related to Fig 5. **A)** Three variable features displayed as color-scale values on the UMAP shown in Fig 5A – AP width appears to vary along UMAP1, while threshold voltage and the depth of the fast AHP additionally distinguish striatal and cortical fast spikers along the UMAP2 axis. **B)** Other significant ion channel-related gene expression differences between striatal and cortical FS interneurons. **C)** Other significant electrophysiological differences between striatal and cortical FS interneurons. **D)** Additional features contrasting PTHLH and TAC3 interneurons. **E)** Differences in morphological feature reflecting the average ratio of Euclidean distance to path distance (tortuosity) between PTHLH and other interneurons. **F)** Additional ion channel-related expression differences that may contribute to intrinsic properties of TAC3 interneurons as compared to other related interneurons. **G)** Example of spontaneous activity and slow plateau-like potentials in SST-CHODL neurons. **H)** Response to step current injections (left, same scale as in G) and comparison of spike width (right) in a LAMP5-LHX8 neuron. This neuron had a calculated input resistance of 147 MOhms and capacitance of 55 pF.

**Figure S7.**
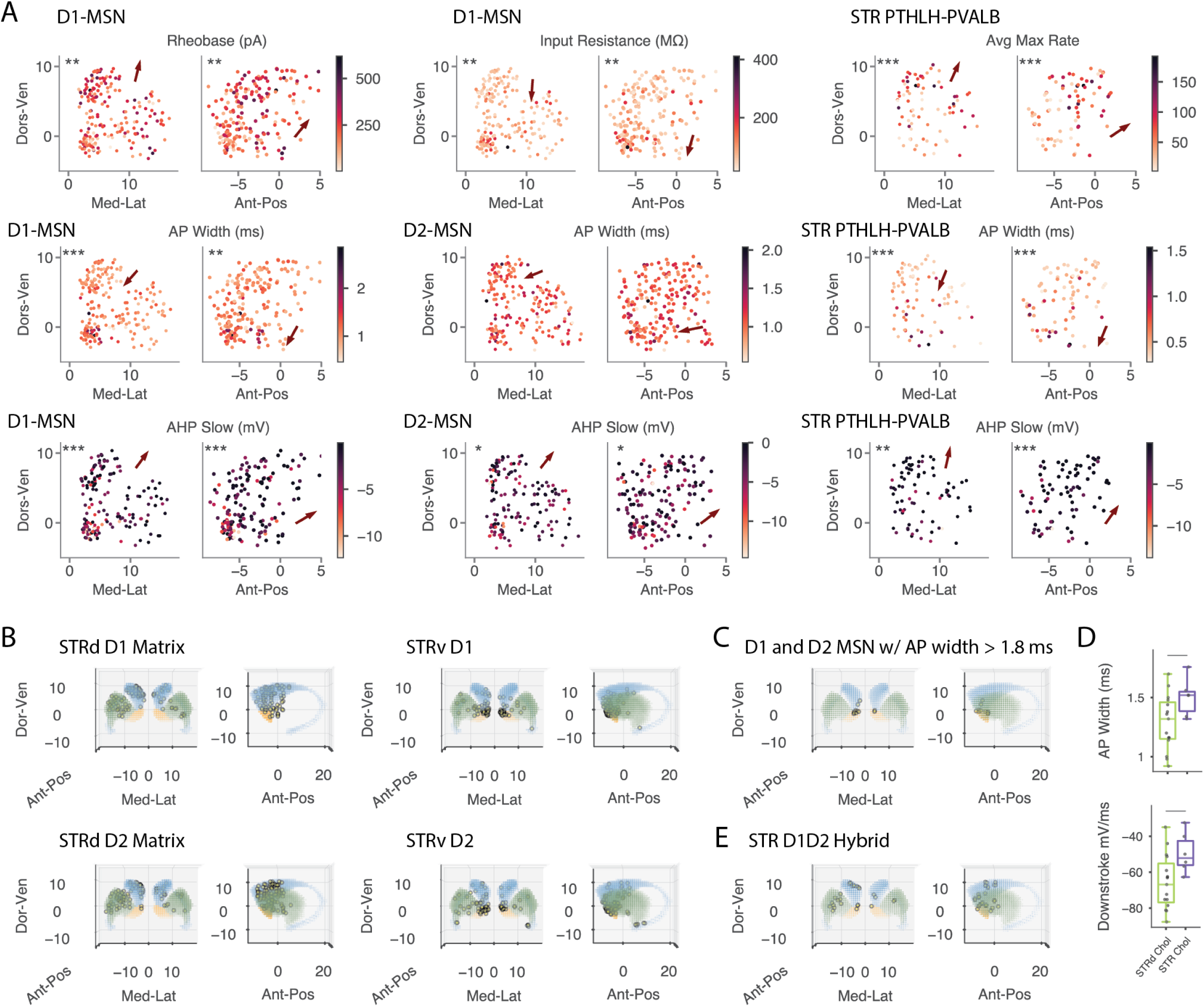
Extended exploration of spatial variation of electrophysiological properties, relating to Fig. 6. **A)** Some features were only significantly varying in MSNs or PTHLH but not both (top row). Other features showed coordinated spatial variation for MSNs and PTHLH, in addition to the ones shown in Fig. 6E. Striosomal cells were excluded from the AHP_slow plot and analysis as they were substantially different for this feature. **B)** Pinned soma locations of dorsal and ventral MSN transcriptomic types were distributed as expected, although ventral transcriptomic mappings were not exclusively localized to ventral structures. Cells in the ventral tail of the caudate appear to map to ventral types.**C)** Locations of MSNs with wide action potentials are plotted in isolation for visibility – the majority localize to ventral striatum. **D)** Although data for cholinergic interneurons was insufficient to do a full spatial analysis, it appears that similar to MSNs and PTHLH neurons, ventral cholinergic neurons had wider spikes and slower AP kinetics. **E)** STR D1D2 Hybrid samples could be found throughout, including in dorsal and ventral striatum.

**Figure S8.**
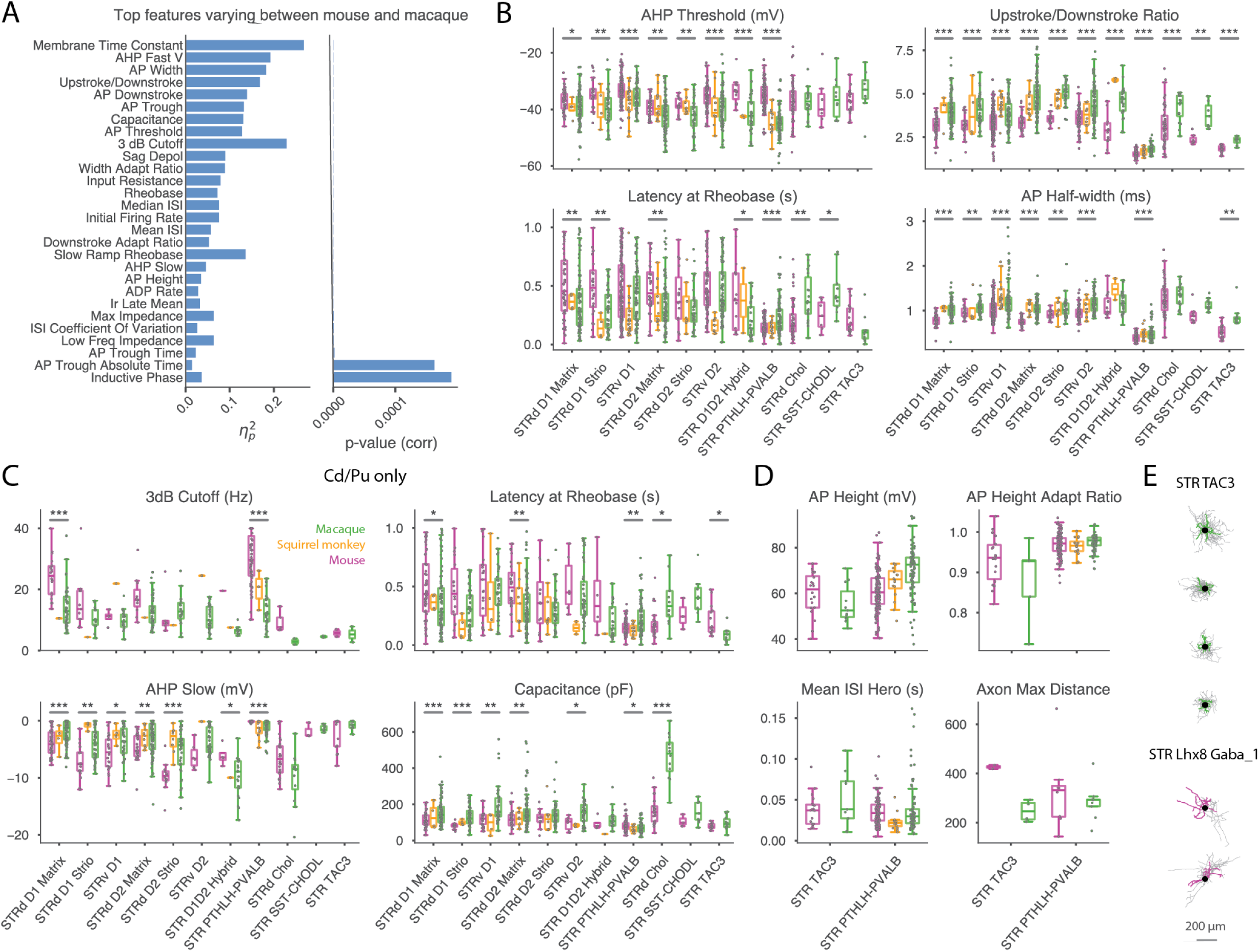
Supporting information on cross-species differences, related to Fig. 7. **A)** Features differing between species in a two-way ANOVA of species and transcriptomic Group, sorted by ascending p-value. This is an extended version of Fig. 7B and shows associated p-values. **B)** Electrophysiological features showing broad systematic variation between mouse and macaque, in addition to those show in Fig. 7C. **C)** Comparisons as in other figures, but restricted to caudate and putamen samples to control for potential confounds arising from species-specific spatial distributions. Although the significance may be diminished due to fewer samples, the conclusions supported remain the same. **D)** Macaque TAC3 IN’s have shorter AP’s than PTHLH IN’s (and all other interneurons, not shown) and strong AP height accommodation, but this is not seen for mouse equivalent types. Mean interspike interval at hero (30-40 pA above threshold) appears similar across groups and species and does not explain differences in last spike time. Mouse STR Lhx8 Gaba_1 (TAC3 equivalent type) axonal arbors may be more extensive than primate TAC3 arbors. **E)** All available morphologies for STR TAC3 and STR Lhx8_GABA_1.

